# SARS-CoV-2 infection of African green monkeys results in mild respiratory disease discernible by PET/CT imaging and prolonged shedding of infectious virus from both respiratory and gastrointestinal tracts

**DOI:** 10.1101/2020.06.20.137687

**Authors:** Amy L. Hartman, Sham Nambulli, Cynthia M. McMillen, Alexander G. White, Natasha L. Tilston-Lunel, Joseph R. Albe, Emily Cottle, Matthew Dunn, L. James Frye, Theron H. Gilliland, Emily L. Olsen, Katherine J. O’Malley, Madeline M. Schwarz, Jaime A. Tomko, Reagan C. Walker, Mengying Xia, Matthew S. Hartman, Edwin Klein, Charles A. Scanga, JoAnne L. Flynn, William B. Klimstra, Anita K. McElroy, Douglas S. Reed, W. Paul Duprex

## Abstract

Vaccines are urgently needed to combat the global coronavirus disease 2019 (COVID-19) pandemic, and testing of candidate vaccines in an appropriate non-human primate (NHP) model is a critical step in the process. Infection of African green monkeys (AGM) with a low passage human isolate of SARS-CoV-2 by aerosol or mucosal exposure resulted in mild clinical infection with a transient decrease in lung tidal volume. Imaging with human clinical-grade ^18^F-fluoro-2-deoxy-D-glucose positron emission tomography (^18^F-FDG PET) co-registered with computed tomography (CT) revealed pulmonary lesions at 4 days post-infection (dpi) that resolved over time. Infectious virus was shed from both respiratory and gastrointestinal (GI) tracts in all animals in a biphasic manner, first between 2-7 dpi followed by a recrudescence at 14-21 dpi. Viral RNA (vRNA) was found throughout both respiratory and gastrointestinal systems at necropsy with higher levels of vRNA found within the GI tract tissues. All animals seroconverted simultaneously for IgM and IgG, which has also been documented in human COVID-19 cases. Young AGM represent an excellent species to study mild/subclinical COVID-19 disease and have shed light on unknown aspects of long-term virus shedding. They are ideally suited for preclinical evaluation of candidate vaccines and therapeutic interventions.

**One Sentence Summary:** Subclinical infection of African green monkeys infected with SARS-CoV-2 results in prolonged shedding of infectious virus from both respiratory and gastrointestinal tracts.

## Main Text

The unprecedented and rapidly spreading coronavirus disease 2019 (COVID-19) pandemic caused by the emerging coronavirus SARS-CoV-2 calls for swift testing of vaccine candidates prior to initiation of human clinical trials. Safety must be prioritized over speed when considering a vaccine that will likely be administered to hundreds of millions of people. Non-human primates (NHPs) serve a unique and important purpose for pre-clinical testing of candidate human vaccines, given their close genetic relatedness with humans. As such, studies in NHPs are geared towards identifying the most appropriate species that recapitulates human disease. Comprehensive dissection of the longitudinal viral pathogenesis in NHP models is critical for successful evaluation of the efficacy of vaccines, antibody therapeutics, and small molecules in pre-clinical studies.

Several NHP models have been reported with SARS-CoV-2 using rhesus and/or cynomolgus macaques and African green monkeys (AGMs) (*1-4*). While used less frequently in biomedical research than rhesus or cynomolgus macaques, AGMs are an old-world NHP species and a natural host for simian immunodeficiency virus (SIV) (*5*). In addition to SIV, AGMs have been used as models for infectious diseases such as Rift Valley fever, pneumonic plague, human parainfluenza virus, SARS-CoV-1, and Nipah virus (*6-10*). To fully understand the pathogenesis of human isolates of SARS-CoV-2 in AGM, it is critical to determine if there are differences between aerosol and mucosal exposure routes, to assess whether clinical-grade imaging of NHPs can be used to detect subclinical infections, and to understand the real-time dynamics of virus shedding.

Young adult male AGMs were infected with a low-passage clinical isolate of SARS-CoV-2 and comprehensive longitudinal disease parameters were compared after either small particle aerosol or multi-route mucosal/intratracheal infection. All AGMs developed mild disease regardless of the route of exposure or infectious dose. Pulmonary lesions were detectable by PET/CT in the acute phase and subsequently resolved. All AGMs exhibited prolonged shedding of infectious virus from oral, nasal, conjunctival, and rectal mucosal surfaces. Viral RNA (vRNA) remained detectable throughout both the respiratory and GI tissues at necropsy in the absence of replication-competent virus. These results show multiple routes are involved in viral shedding and that shedding occurs over a protracted period of time in subclinical animals, both of which will greatly impact SARS-CoV-2 transmission from COVID-19 patients.

### Results: Infection of young AGMs with SARS-CoV-2 resulted in mild respiratory disease

Healthy male AGMs (∼3.5 years of age) were infected with passage 3 (P3) of a human isolate of SARS-CoV-2 from Munich, Germany (Table S1)(*11*). Four animals were infected using a small particle aerosol (designated A1-A4) and two were infected by a multi-route mucosal exposure (designated M1, M2) involving administration of virus into the oral, nasal, and ocular mucosal surfaces and intratracheal instillation using a bronchoscope. Aerosol inhaled exposure doses ranged from 3.7–4.2 log_10_ pfu of virus due to the standard ∼2 log loss of virus after nebulization. Multi-route mucosal exposures were 6.4 log_10_ pfu. After infection, animals were anesthetized for blood draws, mucosal swabs, plethysmography, and chest radiography at regular intervals post-infection.

Clinical disease was mild in all six animals (Fig. S1A). Respiratory function revealed a transient decrease in the volume of air inhaled, or tidal volume, at 7 dpi (Fig. S1B), while other parameters including frequency and expiratory time were within normal limits. In the two mucosally infected AGM and in one aerosol infected AGM, there were several short spikes of fever (maximum deviation 2.4-3.4 °C) during the course of infection (Fig. S2). For these three animals, the average significant elevation in temperature was less than 1 °C for either route of infection, suggesting an overall low-grade fever. The other three aerosol-infected AGMs, A1-A3, developed mild hypothermia response around 5-7 dpi that persisted for most of the remainder of the study (Fig. S2). Complete blood counts (CBC) revealed a transient decrease in lymphocytes and platelets and an increase in neutrophils (Fig. S1); this is also seen in human COVID-19 patients (*12, 13*). Blood chemistry analysis demonstrated decreases in amylase and blood urea nitrogen (BUN) and no elevation in liver enzyme levels (Fig. S3).

### SARS-CoV-2 infected AGMs shed infectious virus from respiratory and gastrointestinal tracts

Virus isolations and q-RT-PCR analysis were performed for oral, rectal, nasal, and conjunctival swab samples obtained over the course of the infection. SARS-CoV-2 isolations were confirmed using indirect immunofluorescence using an anti-spike antibody (Fig. 1A). Syncytia were present *in vitro* isolations, particularly at the later time points (e.g. 21 dpi). Replicating virus was detected in all nasal and oral swabs from all animals on 2 and 4 dpi (Fig. 1B). On 2 dpi, 5/6 rectal swabs and 2/6 ocular swabs were positive for infectious virus. All swabs were broadly negative from all animals at 11 dpi. However, there was a resurgence in the presence of replication competent virus on 14 and 21 dpi in samples from the respiratory and GI tracts (Fig. 1A and B). q-RT-PCR results confirmed the virus isolation results, with a peak in vRNA detection in oral, nasal and conjunctival swabs between 2-7 dpi and a second spike in rectal samples 21 dpi (Fig 1C-F). There was consistent detection of vRNA through 28 dpi although at the stage infectious virus was not isolated from either swabs or necropsy tissues.

**Fig. 1.**
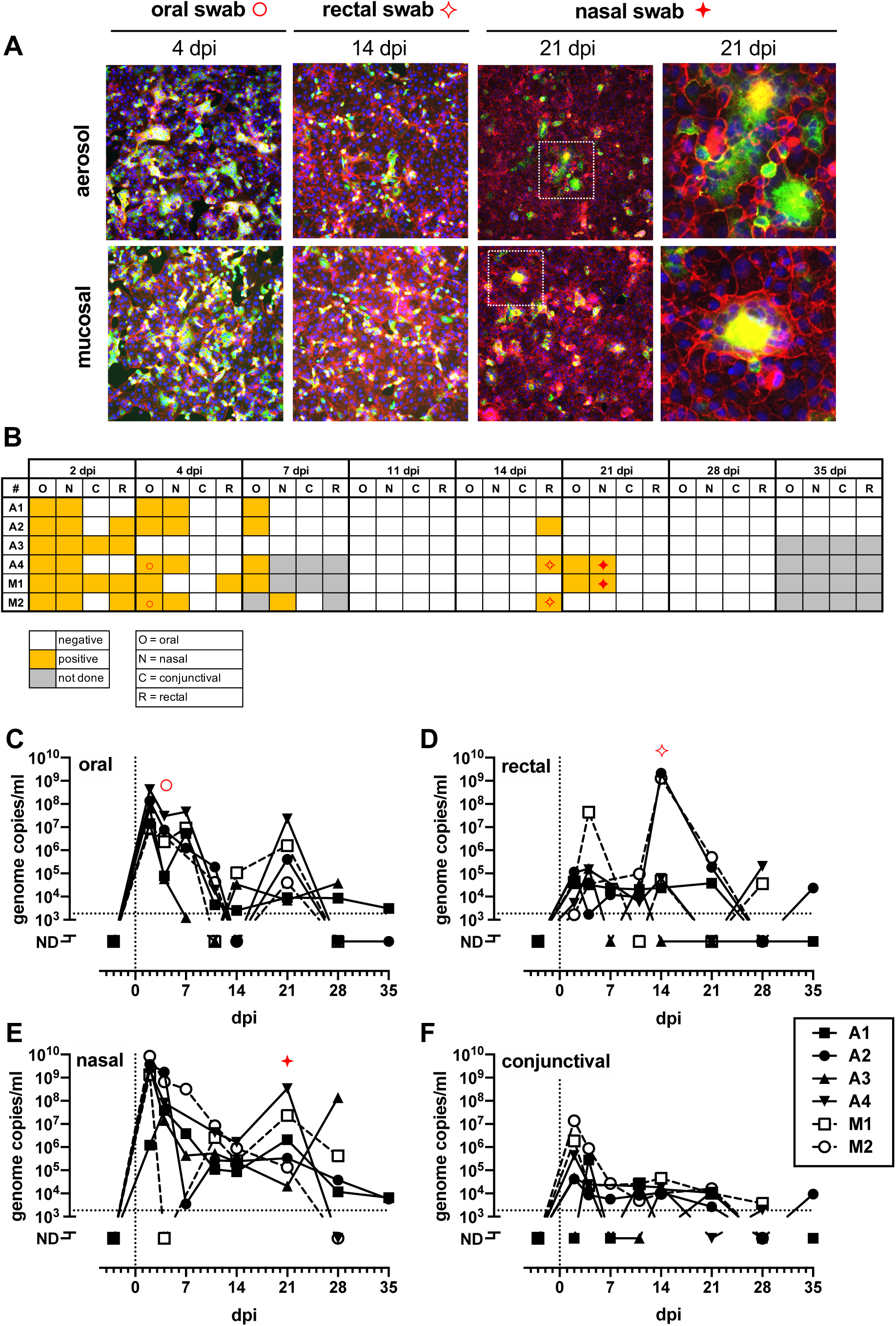
Detection and isolation of SARS-CoV-2 in swabs. (A) Representative virus isolations confirmed by immunofluorescence using anti-SARS2 spike Ab (green) with phalloidin (red) as background and nuclei stained with DAPI (blue). (B) Isolation results from swabs. (B-D) vRNA in swabs measured by q-RT-PCR. AGM infected by aerosol (closed symbols/solid lines; n=4) or multi-route mucosal (open symbols/dashed lines; n=2). Red symbols highlight representative IFA images shown in (A).

### Pulmonary infection in SARS-CoV-2 infected AGM was detected by PET/CT imaging

Molecular imaging using positron emission tomography (PET) with the radiotracer ^18^F-FDG provides a sensitive measurement of metabolic activity within specific anatomical compartments. When co-registered with computed tomography (CT), PET/CT can provide detailed anatomic structure overlaid with areas of high metabolic activity revealed by the tracer. FDG-based PET/CT has been used in NHPs for measurement of lung granulomas caused by infection with *Mycobacteria tuberculosis* (*14-19*). To determine whether lung infection with SARS-CoV-2 could be visualized using FDG-mediated PET/CT or CT alone, imaging was performed pre-infection, 4 dpi, and 11 dpi for most animals. To quantify overall disease burden in the lungs at the various time points, a disease-associated total lung FDG activity level was calculated to measure the total lung inflammation. Analysis of the maximum SUVs in thoracic lymph nodes (LNs) was also conducted.

Both mucosally-infected animals and one of the aerosol animals (A4) had significant lung inflammation at 4 dpi based on FDG uptake; these lesions resolved by 11 dpi (Fig. 2A). The LNs in these three animals also showed the highest FDG uptake at 4 dpi, while all 6 AGM had substantial FDG uptake in the LNs at either 4 or 11 dpi (Fig. 2B). Overall, the extent of disease visualized within the lungs of infected AGMs was modest at all the time points examined (Figs. S4,S5). At 4 dpi, aerosol-infected animal A4 developed areas of disease with a pneumonia appearance in the anterior portions of the accessory lobe and left lower lobe as well as pleural ground glass opacities in the anterior portions of the right and left middle lobes (Fig. 2C). These lesions were resolving by 11 dpi. Peak uptake in the thoracic lymph nodes of this animal was seen at 4 dpi (Fig. 2B). AGM M1 was infected via mucosal exposure and had the most extensive lung disease of all AGMs based on PET imaging. Lesions consisted of diffuse ground glass opacities with corresponding high FDG uptake in the left lower and right upper lobes (Fig. 2D). These areas resolved by day 11, but a new focal area of disease was present that day (cyan arrow; Fig. 2D). Chest radiographs were taken on all longitudinal sampling days. Only animal M1 had detectable radiographic abnormalities with mild non-specific infiltrates present in the left lower lobe at 2 and 4 dpi that were resolving by 7 dpi (Fig S6).

**Fig. 2:**
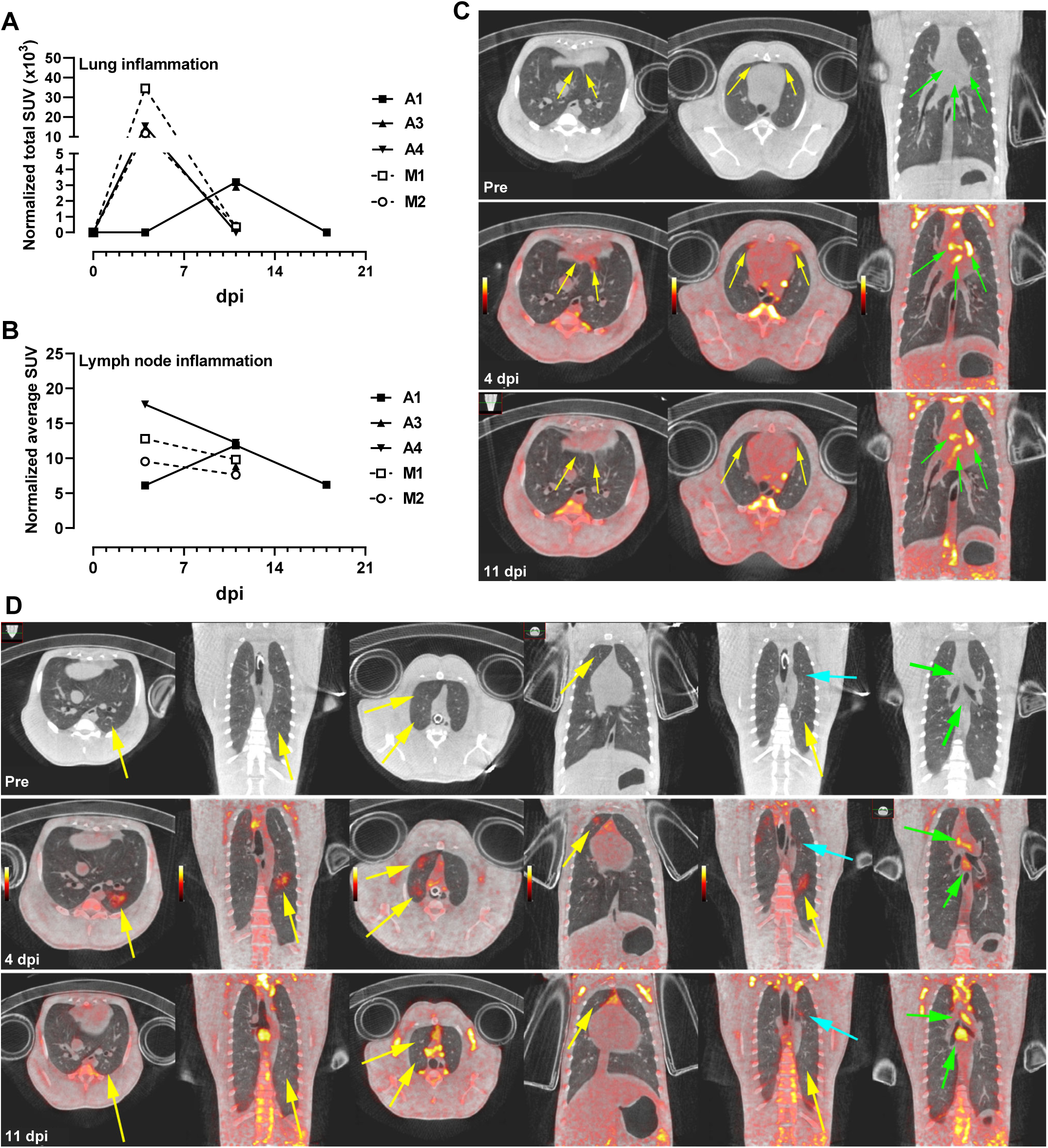
PET/CT imaging of SARS-CoV-2-infected AGM. (A) and (B) Region-of-interest analysis on PET images of animals infected with SARS-CoV-2. (A) Measurement of total lung inflammation via FDG uptake over time. (B) Measurement of average lymph node inflammation over time. Aerosol (closed symbols/solid lines; n=3); multi-route mucosal (open symbols/dashed lines; n=2). Animal A2 is not included in (A) and (B) because only CT scans were performed. (C) For AGM A4 (aerosol), only CT scan was obtained pre-infection; PET/CT at was obtained at 4 dpi and 11 dpi. (D) For AGM M1 (multi-route mucosal), only CT scan was obtained pre-infection; PET/CT at was obtained at 4 dpi and 11 dpi.. Pulmonary infection (yellow arrows); thoracic lymph nodes (green arrows). Cyan arrow highlights new focal area of disease visible on 11 dpi. PET color scale is from 0 - 15 SUV.

### Synchronous seroconversion in SARS-CoV-2-infected AGM

In order to evaluate the kinetics of seroconversion, serial plasma samples from each SARS-CoV-2 infected AGM were assayed for antibodies against the SARS-CoV-2 spike protein receptor binding domain (RBD) using an indirect ELISA (Fig. 3A). All animals generated both IgM and IgG antibodies against SARS-CoV-2. Notably, there was simultaneous seroconversion of both IgM and IgG. This contradicts the classic immunology paradigm of IgM proceeding IgG in response to antigen exposure, however this mirrors precisely what occurs in COVID-19 patients (*20-23*), further validating the utility of this AGM model. All animals seroconverted in the second week post exposure and those that were exposed via the mucosal route had higher overall titers than those that received the virus via the aerosol route. This could reflect the fact that mucosal infected animals received a higher challenge dose than aerosol infected animals. Neutralization of SARS-CoV-2 was measured in matched samples using a plaque-reduction neutralization-80% test (PRNT_80_) (Fig. 3B). Neutralizing antibodies were present in samples collected 7-11 dpi. There were no differences in kinetics or activity between the two exposure routes demonstrating that higher dose received during multi-route mucosal infection neither affected the onset nor strength of the humoral immune response.

**Fig. 3:**
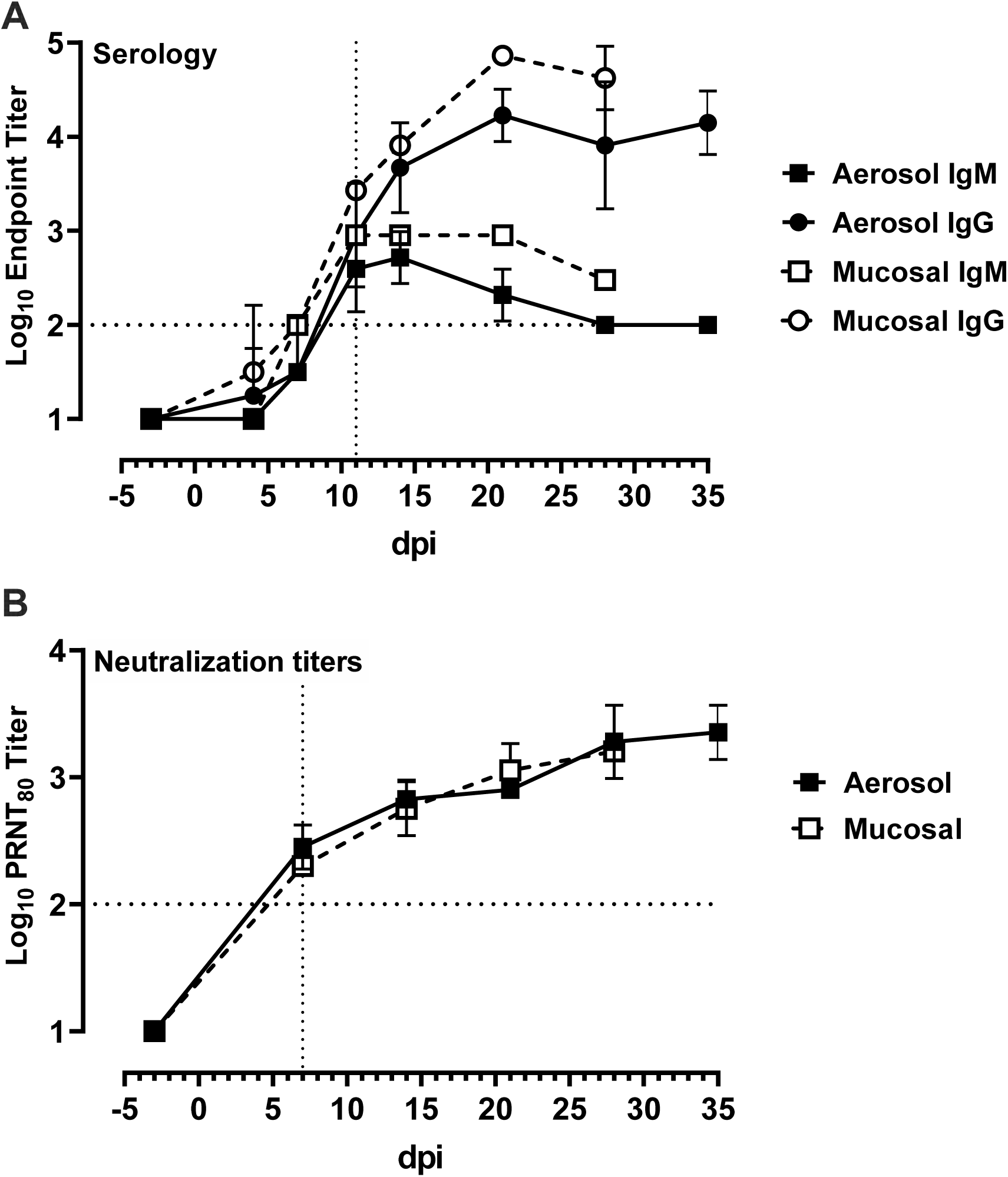
Seroconversion and neutralization in SARS-CoV-2 infected AGMs. Serial plasma samples were assayed for virus specific antibodies or neutralization capacity. (A) virus-specific IgG and IgM were measured using a SARS-CoV-2 spike receptor binding domain-based ELISA. (B) neutralization titers were determined by a plaque-reduction neutralization 80% assay (PRNT_80_). The horizontal dotted line represents the limit of detection of each assay.

### Cytokinemia and immune activation over time revealed early response to viral infection

PBMC were isolated from whole blood and stained for flow cytometry using either a myeloid or lymphoid panel (Table S2; Figs. S7,S8). The proportions of proliferating (Ki-67+) CD4 and CD8 T cells increased transiently between 2-11 dpi in most animals (Fig. S9). Unlike in the aerosol infection, there were sustained increases in proliferating CD8 T cells in the mucosally-infected animals (Fig. S9D) which is similar to what occurs in human COVID-19 patients (*12, 13*). NK cell frequencies varied over the course of the experiment as did the mean fluorescence intensity (MFI) of CD16 on NK cells with no clear pattern emerging (Fig. S9E,F). This is relevant since COVID-19 patients with severe, but not moderate, disease demonstrated decreased CD16 MFI on NK cells (*12*). Most notably, an increased frequency of CD20^lo^ B cells was noted in all animals in the second week of infection, accompanied by increased expression of Ki-67 in these cells, which is consisted with a plasmablast phenotype (Fig. S9H,I). In comparison, CD20^hi^ B cells saw no increase in Ki-67. An NHP-specific multiplex cytokine and chemokine assay was used to measure the levels of 30 analytes in longitudinal plasma samples. The majority of cytokines, including IFN-α, IFN-γ, IL-6, and TNF-α, were undetectable in all animals at all time points tested. For the cytokines that were detectable, most AGMs had early transient peaks in expression of MCP-1, IL-1RA, IP-10, and ITAC at 2 dpi (Fig. S11), indicative of a general response to viral infection. Peak plasmablast frequency (Fig. S9H) and B lymphocyte chemoattractant (BLC) expression (Fig. S11E) coincided with the appearance of virus specific antibodies (Fig. 3).

### SARS-CoV-2 vRNA was detected throughout respiratory and gastrointestinal tract at necropsy

Upon euthanasia of each animal at 28 dpi (35 dpi for A1 and A2), full necropsies were performed and tissues were tested by qRT-PCR for levels of vRNA (Fig. 4). vRNA was widespread throughout the upper and lower respiratory tracts including the soft palate and nasal turbinates. The overall levels of vRNA detected were comparable between the two exposure routes and doses, indicating that the 100-fold higher infectious dose that the mucosally infected animals received did not translate into higher levels of tissue vRNA at necropsy. More vRNA was detected along the entire gastrointestinal (GI) tract compared to the lower and upper respiratory tracts, highlighting differential tropism of SARS-CoV-2. vRNA was detected in 5/6 olfactory bulbs but was largely absent from the cortex and cerebellum of most animals. The heart, liver, spleen, and kidney were largely devoid of vRNA. Virus isolation attempts on lung and gastrointestinal tissue obtained at necropsy were unsuccessful.

**Fig. 4.**
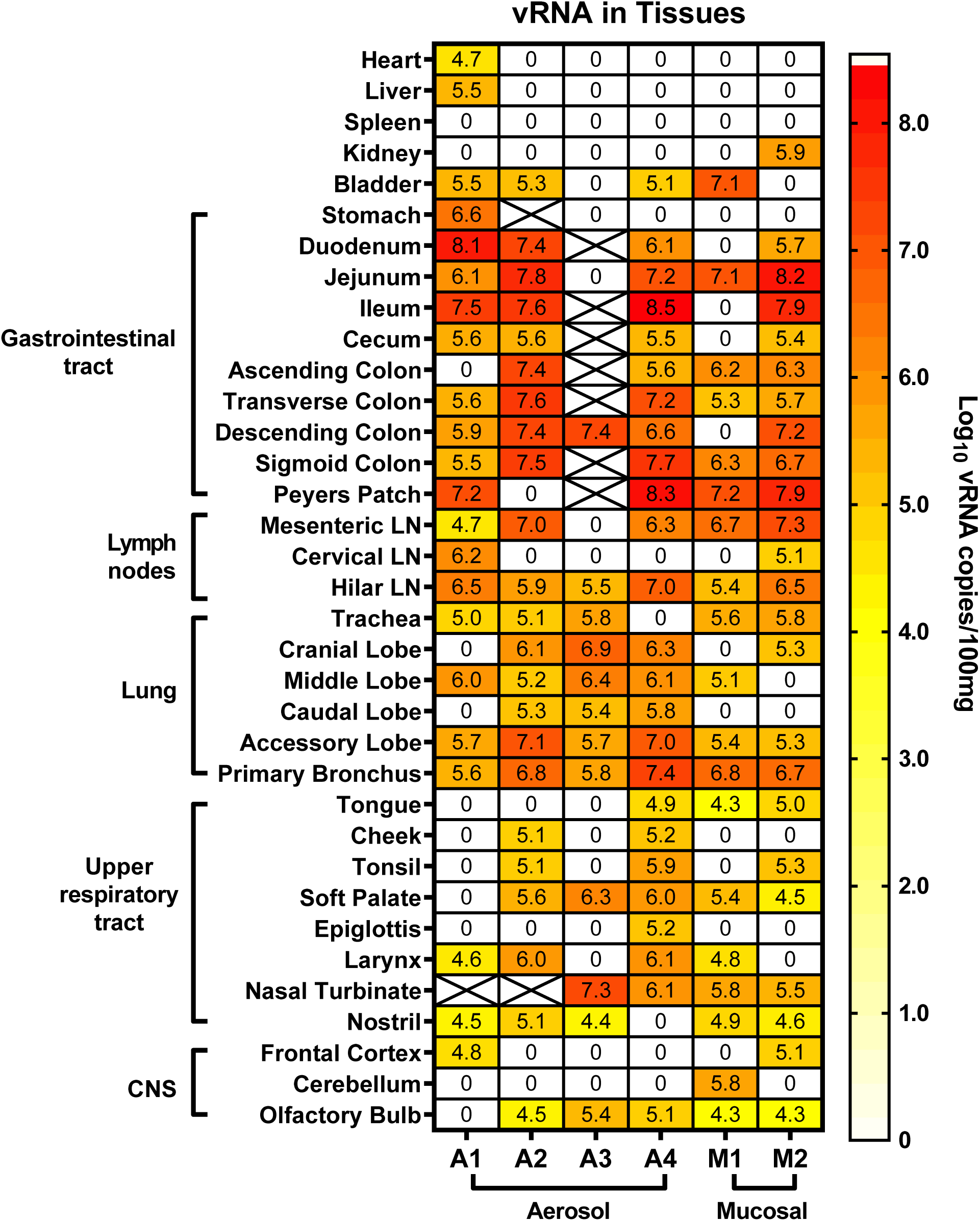
Viral RNA within tissues of SARS-CoV-2 infected AGM. Animals underwent necropsy at 28 dpi (35 dpi for A1 and A2) and the indicated tissues were extracted and tested for viral RNA by q-RT-PCR. Heat map shows the log-transformed vRNA copies per 100 mg of tissue. X indicates the sample was not available or not tested.

### Residual histopathology detected at necropsy

Since the AGMs were euthanized at 28 or 35 dpi, tissue specimens for acute histopathology were not obtained. However, lung samples and lymph nodes from animals known to be positive by PET/CT were taken at necropsy and examined by a board-certified veterinary pathologist. Importantly, even 4-5 weeks after infection, several animals demonstrated multiple pulmonary foci of interstitial infiltration and expansion by either lymphocytic or mixed inflammatory infiltrates (Fig. 5). While not a specific diagnostic finding, this is a common pathological effect of many pneumotropic viruses and is also consistent with the convalescent stage of a respiratory infection. Interestingly, multiple syncytiated cells were observed in the germinal follicles of Peyer’s patches from two animals (Fig 5B). Given the frequent propensity of coronaviruses, including SARS-CoV-1, to lead to fused cells *in vitro*, our observation that isolated virus from the AGMs form syncytia in Vero-E6 cells (Fig. 1A) and the observation of multinucleated syncytial cells in human cases of disease (*24, 25*), this finding is likely a viral cytopathic effect. Moreover, the Peyer’s patches from 5 out of 6 AGMs contained some of the highest levels of vRNA (Fig. 4).

**Fig. 5.**
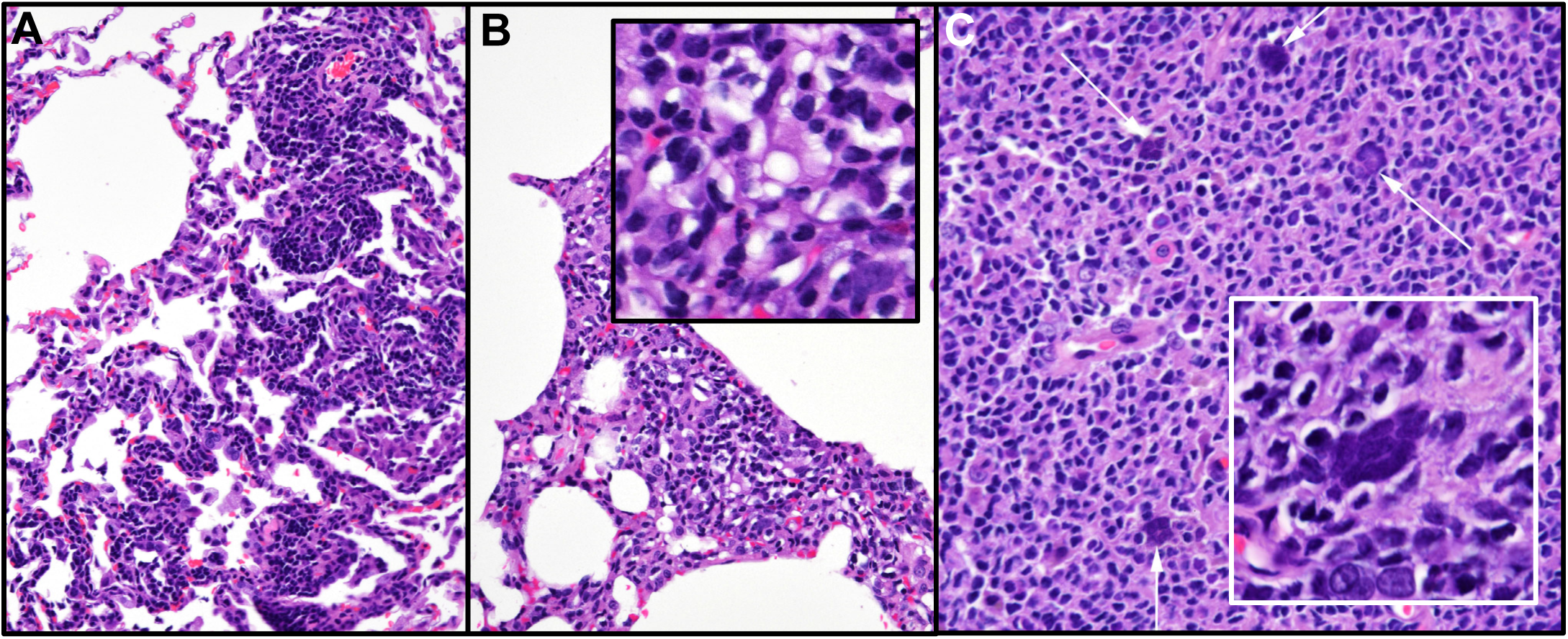
Histopathology detected at necropsy. (A) and (B) Lung (20X) reveals pulmonary foci of mild infiltration and interstitial expansion by lymphocytic and mixed inflammatory cells. (C) Peyer’s patch (20X) shows multiple, syncytialized cells (white arrows). Inset of Peyer’s patch illustrates smudgy syncytia. Histopathology interpreted by a board-certified veterinary pathologist.

## Discussion

AGMs were used as a non-human primate model for SARS-CoV-1 infection after the 2003 epidemic. Compared to rhesus macaques, AGMs supported enhanced SARS-CoV-1 virus replication and developed more severe lung pathology (*10, 26*). They are also an excellent species to model other respiratory diseases such as pneumonic plague and human parainfluenza virus (*7, 9*). In this study, COVID-19 disease in young AGMs was mild, reflecting what frequently occurs in young, healthy humans infected with SARS-CoV-2 (*27, 28*). Low-grade fever was observed and respiratory symptoms were limited to a transient decrease in tidal volume. Advanced imaging using a metabolic PET probe permitted visualization of pulmonary lesions undetectable by other modalities such as standard chest radiographs. PET/CT is used clinically in managing COVID-19 patients and is effective at visualizing lesions in asymptomatic patients (*29, 30*). It is notable that the three AGM (A4, M1, M2) that had the most febrile responses also had the highest level of inflammation in the lungs and lymph nodes as detected by PET/CT. PET/CT imaging was also performed in crab-eating macaques infected intra-tracheally with SARS-CoV-2 (*31*). Disease in the macaques was mild/subclinical, and PET/CT scans showed areas of pneumonia and consolidation that were FDG avid. The importance of these studies lies in demonstrating that PET/CT imaging can be used to bridge human animal data, particularly with respect to determining reductions in disease burden following vaccination or therapeutic interventions.

A key finding here was that virus isolations followed by immunofluorescence confirmation demonstrated unequivocally that infectious virus was shed from the respiratory and GI early followed by a recrudescence around 14-21 dpi. The majority of *in vivo* SARS-CoV-2 studies use vRNA detection as a surrogate for infectious virus. The biphasic shedding pattern was unexpected and highlights the need to study longer term infection in animal models. vRNA was still detectable at substantial levels throughout the entire GI tract at 4-5 weeks after infection, and GI titers were 10-100-fold higher than what was found throughout the upper and lower respiratory tracts. This demonstrates the tropism of SARS-CoV-2 for intestinal tissues and the importance of GI replication in the viral pathogenesis. Transmission of SARS-CoV-1 through fecal matter contributed to super-spreader events during the 2003 outbreak (*32, 33*), and the mechanism underlying this remains largely unexplored. For SARS-CoV-2, vRNA and infectious virus has been detected in feces of COVID-19 patients with or without GI symptoms (*34-36*). The AGM model will allow an understanding of how SARS-CoV-2 is shed from the GI tract and potentially transmitted via feces.

In a recent study with cynomolgus macaques, SARS-CoV-2 infection was mild to subclinical even in aged animals, shedding of vRNA from mucosal surfaces was limited to early time points, and vRNA at necropsy was largely limited to respiratory tissues (*1*). A preprint study in rhesus macaques showed transient clinical disease marked by brief fever right after infection, transient weight loss, mild respiratory depression, and pulmonary infiltrates by chest radiograph (*2*). Upon necropsy, vRNA was found primarily in the lungs and upper respiratory tract and not in the GI tract. Another preprint study in rhesus macaques found similar limited disease with shedding of vRNA and infiltrates by chest radiograph (*3*).

The current consensus regardless of NHP species or viral isolates is that infection with SARS-CoV-2 leads to mild or subclinical infection, with shedding of virus from the respiratory tract. Severe disease or lethality has been rarely seen after either mucosal or aerosol exposure. One distinguishing feature of the AGM model appears to be tropism of virus for tissues throughout the entire gastrointestinal tract, as we found here, including substantial shedding of infectious virus. A key finding from this study was the repeated isolation of infectious virus over time from mucosal swabs including the rectum. This is critically important for the fundamental understanding of both COVID-19 disease and person-to-person transmission.

The SARS-CoV-2-specific immune responses occurred in the second week of infection. The concurrent detection of both IgM and IgG against the receptor binding domain of the spike protein by ELISA coincided with the appearance of a neutralizing antibody response. Mucosally-infected animals had higher ELISA titers than aerosol exposed animals and increased levels of the B cell chemokine CXCL-13 in the plasma during the first week preceding the seroconversion. Plasmablasts peaked in all infected animals on 10 dpi. While a greater magnitude of B cell response seen in mucosal animals could have been secondary to higher inoculum dose, it could also reflect exposure route of which could have implications for optimal immunization. Virus specific T cell responses were not assessed directly. However, proliferation of both CD4 and CD8 T cells, as evidenced by Ki-67 staining, occurred in all animals in the first two weeks of infection. These data are analogous to what occurs in humans with moderate COVID-19 disease (*12, 13*).

The source of NHP used in pathogenesis and efficacy studies can play an important role in clinical outcome, and thus clear explanation of the source of animals and genetic background for future studies is important. The AGMs used in our study were from the Vervet Research Colony at Wake Forest University. The colony founders were 57 wild-caught animals from St. Kitts in the 1970’s (*37, 38*). The animals used here were all male and all ∼3.5 years of age (age range is 25-35 years in captivity), making them young and around the age of sexual maturity. Future studies should consider using older animals and/or those directly imported from St. Kitts. Particularly relevant to the COVID-19 pandemic is that AGM from the West Indies spontaneously develop hypertension without dietary intervention or inbreeding (*39*). Moreover, they naturally develop type II diabetes and atherosclerosis, making them an excellent species to study pathogenesis of SARS-CoV-2 under conditions representative of the comorbidities associated with severe COVID-19 in humans (*40, 41*). Increasing age, genetic variability, and spontaneous hypertension, diabetes, and cardiovascular disease may lead to more severe outcomes in AGMs when infected with SARS-CoV-2, and thus may more accurately reflect the variation seen in the human population.

In summary, we comprehensively evaluated the pathogenesis of SARS-CoV-2 in the AGM model. Our study suggests that AGMs are excellent models for many aspects of COVID-19 in humans: 1) young healthy AGMs may represent subclinical or mild human disease, 2) consistent shedding of infectious virus from respiratory and GI tracts in the absence of overt disease allows experimental understanding of the mechanisms underlying tissue tropism and transmission, and 3) PET/CT imaging modalities are effective at detecting SARS-CoV-2 infection. Critical vaccine trials could use the AGM model to measure virus shedding and/or lung lesions by PET/CT after post-vaccination challenge as measures of vaccine efficacy.

## Acknowledgments

We thank Dr. Natalie Thornburg (CDC Atlanta) and Drs. Christen Drosten and Jan Felix Drexler (Charité – Universitätsmedizin Berlin) for providing SARS-CoV-2 isolates. We also thank Dr. Carin Beth Ahner, DVM and Stacey Barrick for critical assistance with the animal studies. Dr. Seema Lakdawala generously provided SARS-CoV-2 spike protein for serological assays. We appreciate the efforts of the University of Pittsburgh Department of Environmental Health and Safety for timely regulatory and logistical oversight this work under pandemic conditions.

## Funding

Funding for this study was generously provided by Dr. Patrick Gallagher, Chancellor of the University of Pittsburgh and by the Center for Vaccine Research. Funding for the PET/CT Reading Core was through the Bill and Melinda Gates Foundation (OPP1130668). A.M. was funded through a Burroughs Wellcome career award for medical scientists (award 1013362) C.M was funded through NIH T32 AI060525.

## Author contributions

Conceptualization: W.P.D., A.H., D.R., W.K., A.M.; Investigation: A.H., S.N., C.M., A.W., N.T-L., J.A., E.C., M.D., L.J.F., T.G., E.O., K.O., M.S., J.T., R.W., M.X., A.M., D.R. Formal analysis: A.H., A.W., M.H., E.K., C.S., J.F., D.R., W.K., A.M., W.P.D.; Writing (original draft): A.H.; Writing (review and editing): W.P.D., A.H., D.R., W.K., A.M., C.S., J.F.; Visualization: W.P.D., A.H., D.R., W.K., A.M., A.W., C.S., J.F.; Project administration: W.P.D., A.H., D.R., W.K., A.M. Funding acquisition: W.P.D.

## Competing interests

The authors declare no competing interests.

## Data and materials availability

All data are available in the main text or supplementary material.

## Supplementary Materials

Materials and Methods

Figures S1-S11

Tables S1-S2

References (*1-6*)

## Notes

### Competing Interest Statement

The authors have declared no competing interest.

