## Supplementary Figures for "SARS-CoV-2 infection of African green monkeys results in mild respiratory disease discernible by PET/CT imaging and prolonged shedding of infectious virus from both respiratory and gastrointestinal tracts"

**This PDF file includes:**

Materials and Methods
Figs. S1 to S11
Tables S1 to S2
References 1-6

### Materials and Methods

**Ethics.** The animal work performed adhered to the highest level of humane animal care standards. The University of Pittsburgh is fully accredited by the Association for Assessment and Accreditation of Laboratory Animal Care (AAALAC). All animal work was performed under the standards of the Guide for the Care and Use of Laboratory Animals published by the National Institutes of Health (NIH) and according to the Animal Welfare Act guidelines. All animal studies adhered to the principles stated in the Public Health Services Policy on Humane Care and Use of Laboratory Animals. The University of Pittsburgh Institutional Animal Care and Use Committee (IACUC) approved and oversaw the animal protocols for these studies.

**Biological safety.** All work with SARS-CoV-2 was conducted under biosafety level-3 (BSL-3) conditions in the University of Pittsburgh Center for Vaccine Research (CVR) and the Regional Biocontainment Laboratory (RBL). Respiratory protection for all personnel when handling infectious samples or working with animals was provided by powered air-purifying respirators (PAPRs; Versaflo TR-300; 3M, St. Paul, MN). Liquid and surface disinfection was performed using Peroxigard disinfectant (1:16 dilution), while solid wastes, caging, and animal wastes were steam sterilized in an autoclave.

**Virology**. The SARS-CoV-2 isolate used was a passage 3 (p3) of the Munich isolate described previously (*1*). Virus was titrated by plaque assay and titers are expressed as plaque forming units (pfu), and infection was visualized in cells by indirect immunofluorescence, as described (*1*).

**General animal procedures**. Six male AGMs were obtained from the Vervet Research colony at Wake Forest University (Table S1). All were captive bred animals originally from St. Kitts (*2*). They were serologically negative for herpes B virus, SIV, simian T-cell leukemia virus (STLV), and simian retrovirus (SRV). During quarantine, animals underwent telemetry implant surgery, described below. For euthanasia, each animal was sedated with 20 mg/kg ketamine, followed by injection of 200 mg/kg of Beuthanasia IV. Following euthanasia, each animal was perfused via the left ventricle with saline using a variable perfusion machine (GP1000; Fisher Scientific).

**Telemetry surgery and data acquisition**. Each AGM was implanted with a DSI PhysioTel Digital radiotelemetry transmitter (DSI Model No. M00) capable of continuously recording body temperature. A subcutaneous pocket was created on the left lateral aspect of the abdomen, and the telemetry implant was placed in the pocket and closed using skin sutures. During acquisition, data was transmitted from the implant to a TRX-1 receiver mounted in the room connected via a Communications Link Controller (CLC) to a computer running Ponemah v6.5 (DSI) software. Pre-exposure data collection began at least seven days in advance of infection. Data collected from Ponemah was exported as 15-minute averages into Excel files which were subsequently analyzed in MatLab 2019a. Using pre-exposure baseline data, an auto-regressive integrated moving average (ARIMA) model was used to forecast body temperature assuming diurnal variation across a 24-hour period. The code is available at https://github.com/ReedLabatPitt/Reed-Lab-Code-Library. Residual temperatures were calculated as actual minus predicted temperatures. Upper and lower limits to determine significant changes were calculated as the product of 3 times the square root of the residual sum of squares from the baseline data.

**Aerosol infection.** Aerosol exposures were performed using the Aero3G aerosol management platform as previously described (*3*). Jacketed External Telemetry Respiratory Inductive Plethysmography (JET-RIP; DSI) belts were placed around the abdomen and chest of the animal and calibrated to a pneumotach. This allowed monitoring and recording of respiratory function including minute volume during the exposure via the Ponemah v5.4 software platform (DSI). Aerosols were generated using an Aerogen Solo vibrating mesh nebulizer as previously described (*4*) with a total airflow of 16 l/min into the chamber. Aerosol sampling was performed with an all-glass impinger (AGI) operating at 6 l/min, -6 to -15 psi. Particle size was measured once during each exposure at 5 minutes using an Aerodynamic Particle Sizer (TSI, Shoreview, MN). Inhaled dose was calculated based on pre- and post-sampling titers as described (*3*).

**Multi-route mucosal infection**. Virus was administered to two AGM (5 ml of $5x10^5$ pfu/ml) split between 4 sites (nasal (0.5 ml per nare), oral (1 ml), ocular (100 µl, tracheal (2.8 ml)). Intra-tracheal infections were performed using a Wolf bronchoscope; the distal end of the scope was passed between the laryngeal folds into the trachea until the bifurcation of the trachea was visualized. Once the scope was in place, approximately 2.8 ml of virus stock solution was administered into the trachea, followed by 3 ml of sterile saline, and 5 ml of air to flush any remaining fluid from the bronchoscope into the airway.

**Clinical Scoring.** Clinical signs were recorded twice daily and each animal was given an objective score for each of 3 components. The total score is the sum of all 3 component scores.
Temperature: Normal range (any temp that equals or falls between the upper and lower temperature limits) = 0; significant elevation (above the upper limit) = 1; significant decrease (below the lower limit) = 2; severe hypothermia (<34ºC but >31ºC) = 3; temperature <31ºC = euthanize promptly.
Clinical Appearance/Behavior: normal = 0; lethargic, huddled = 1; piloerection, dehydration, or anorexia = 2; moves only when prodded = 3; no response to prodding = euthanize promptly
Respiratory Signs: none = 0; nasal discharge = 1; increased respiratory rate and effort = 2; respiratory distress (defined as respiratory rate greater than twice the baseline rate) = 3; rales = euthanize promptly.

**Plethysmography**: Respiratory function was assessed in anesthetized animals using a head-out plethysmography chamber and pneumotach connected to a digital preamplifier run by Finepointe v2.8 software (DSI). For purposes of this study, we used a universal study modified to collect data similar to the chronic obstructive pulmonary disease (COPD) studies that Finepointe has established for whole-body plethysmography chambers was used. The chamber, pneumotach, and preamplifier were calibrated before use,data was collected for three minutes and analyzed within Finepointe.

**Longitudinal sampling**. Animals were sedated for sampling after infection by administration of 10 mg/kg ketamine via intramuscular injection. Once sedated, 2-3 ml of blood was drawn from either the right or left femoral vein into an EDTA tube. Plasma was frozen for serologic, immunologic and virologic assays, and peripheral blood mononuclear cells (PBMCs) were prepared as previously described for flow cytometric studies (*3*). CBC analysis was performed using the Abaxis HM5 hematology analyzer. Blood chemistry analysis was performed using the Comprehensive Diagnostic Panel rotor (Abaxis 500-0038) on an Abaxis VS2 chemistry analyzer. Radiographs and mucosal swabs were obtained from each animal. Four types of swabs were obtained at each time point: oral, ocular, nasal, and rectal. Swabs were rotated in place for 10 seconds and then inserted into 1 ml virus transport media (495 mlml Opti-MEM and 5ml of 100 ug/ml Antibiotic/Antimycotic). Each swab was vortexed for 5 seconds, centrifuged to collect the media, and then the media was frozen. For erythrocyte sedimentation rate (ESR), 1 ml of EDTA treated blood was placed into measured capillary tubes for 1 hour. Sedimentation of red blood cells was then recorded in centimeters.

**PBMC thawing and staining**. For flow cytometry, PBMC samples were thawed in a 37°C water bath and resuspended in DMEM:F12 media (Fisher 11320-033). Cells were then pelleted (1000 rpm x 5 min) and washed twice using DPBS. Pellets were resuspended in live/dead solution (Fisher L34961) on ice for 20 minutes. Samples were then washed twice with FACS buffer and stained with extracellular antibodies consisting of CD14 (BD 561391), CD16 (BD 562874 and BD561394), CD11b (BD 561887), HLA-DR (BD 339194), CD3 (BD 558124 or BD 557757), CD20 (BD 560734), CD4 (BD 347327), CD8 (BD 335787), NKG2A (Beckman Coulter A60797), CD1c (Biolegend 331506), and CD123 (BD 560826) on ice for 30 min. Stained samples were washed twice with FACS buffer and fixed and permeabilized using BD Cytofix/Cytoperm (BD554714) for 20 min on ice. Samples were washed twice using 1x BD Cytoperm buffer (BD554714) 500 g x 4 min and resuspended in intracellular antibody cocktail consisting of CD38 (BD 560676) and Ki-67 (BD 558616) for 20 minutes on ice. After washing, samples were fixed and inactivated using 4% (w/v) paraformaldehyde, run on a BD LSRII, and analyzed using FlowJo 10.5.0.

**Multiplex cytokine analysis**. ProcartaPlex NHP Cytokine & Chemokine Panel 30plex (Fisher EPX300-400-44-901) was used. For each sample, 25 µl of plasma was used following manufacturer's instructions. Samples were run on BioRad Bio-Plex 200 reader and results analyzed using Bio-Plex manager software v6.2.

**Radiography.** Using a Sedecal Portable X-ray unit with a Ralco X-ray collimator, each sedated animal was placed in ventral dorsal recumbency on top of a double bagged Fugi (25.2 x 30.3 cm or 35.4 x 43.0 cm) cassette. The collimator was focused on the thorax of the animal. The X-ray unit was set to 55 kvp and 2.0 MAS for 0.07 sec. Radiographs were processed using a Med Serv plus digital processor and were interpreted by a board-certified radiologist.

**PET/CT imaging**. Animals were sedated with 10 mg/kg ketamine / 0.5 ml atropine before imaging. An intravenous catheter was placed in the saphenous vein and animals were injected with ~ 5 millicurie (mCi) of $^{18}$F-FDG. An endotracheal tube was placed for ventilation during scanning and the eyes were lubricated with artificial tears. Once placed on the imaging bed, anesthesia was induced with 2.5 to 3% isoflurane which is reduced to 0.8 – 1.2% for maintenance. Breathing during imaging was maintained using an Inspiration 7i ventilator (eVent Medical, Lake Forest, CA, USA) with the following settings: PF = 9.0 l/min, respiration rate = 18 – 22 bpm, tidal volume = 60 ml, O2 = 100, PEEP = 5 – 8 cm $H_2O$, peak pressure = 15 – 18 cm $H_2O$, I:E ratio = 1:2.0. A breath hold was conducted during the entirety of the CT acquisition.

PET/CT scans were performed on a MultiScan LFER 150 (Mediso Medical Imaging Systems, Budapest, Hungary). CT acquisition was performed using the following parameters: Semi-circular single field-of-view, 360 projections, 80 kVp, 670 µA, exposure time 90 ms, binning 1:4, voxel size of final image: 500 x 500 µm. PET acquisition was performed 55 min after intravenous injection of $^{18}$F-FDG with the following parameters: 10 min acquisition, single field-of-view, 1-9 coincidence mode, 5 ns coincidence time window. PET images were reconstructed with the following parameters: Tera-Tomo 3D reconstruction, 400-600 keV energy window, 1-9 coincidence mode, median filter on, spike filter on, voxel size 0.7 mm, 8 iterations, 9 subsets, scatter correction on, attenuation correction based on CT material map segmentation. Serial CT or PET/CT images were acquired pre-infection and at 4 and 11 dpi. Animal A2 was CT scanned at 9 dpi instead of 11 dpi, and animal A1 was scanned at 18 dpi in addition to the standard imaging schedule previously described.

Images were analyzed using OsiriX MD or 64-bit (v.11, Pixmeo, Geneva, Switzerland). Before analysis, PET images were Gaussian smoothed in OsiriX and smoothing was applied to raw data with a 3 x 3 matrix size and a matrix normalization value of 24. Whole lung FDG uptake was measured by first creating a whole lung region-of-interest (ROI) on the lung in the CT scan by creating a 3D growing region highlighting every voxel in the lungs between -1024 and -500 Hounsfield units. This whole lung ROI was copied and pasted to the PET scan and gaps within the ROI were filled in using a closing ROI brush tool with a structuring element radius of 3. All voxels within the lung ROI with a standard uptake value (SUV) below 1.5 were set to zero and the SUVs of the remaining voxels were summed for a total lung FDG uptake (total inflammation) value. Thoracic lymph nodes were analyzed by measuring the maximum SUV within each lymph node using an oval drawing tool. Both total FDG uptake and lymph node uptake values were normalized to back muscle FDG uptake that was measured by drawing cylinder ROIs on the back muscles adjacent to the spine at the same axial level as the carina (SUVCMR; cylinder-muscle-ratio) (*5*). PET quantification values were organized in Microsoft Excel and graphed using GraphPad Prism.

**Tissue Extraction and Processing.** For whole tissues, 100 mg of tissue was harvested, suspended in 1.5 ml DPBS (no cations) supplemented with 1% (v/v) fetal bovine serum (FBS) and penicillin-streptomycin [100 iU/100 µg/ml], homogenized using an Omni tissue homogenizer (Omni International). Tissue homogenate or swab eluate (100 µl) was added to 900 µl of Tri-Reagent (ThermoFisher), thoroughly mixed by vortexing. To ensure virus inactivation, the samples were incubated for 10 minutes at room temperature, stored overnight at -80 °C prior to removal from the BSL-3 facility. Subsequent storage at -80 °C or RNA isolation and one-step qRT-PCR analyses were performed at BSL-2.

**RNA Isolation and one-step qRT-PCR.** Tissue and swab RNA was isolated using a standard alcohol precipitation method and eluted in 40 µl of nuclease-free water. For swab samples, 5 µl of polyacryl carrier (Molecular Research Center) was added to the specimen/Tri-Reagent mixture and incubated at room temperature for 30 seconds. The RNA isolation procedure described for tissues was followed for the remainder of the isolation. For quantitation of viral RNA, a multiplex one-step qRT-PCR was performed using the 4x Reliance One-Step Multiplex RT-qPCR Supermix (BioRad), as described (*1*), using primer and probes targeting the nucleocapsid (N) designed and optimized by the Centers for Disease Control and Prevention (2019-nCoV_N2 forward primer 5'-TTACAAACATTGGCCGCAAA-3'; 2019-nCoV_N2 reverse primer 5'-GCGCGACATTCCGAAGAA-3' and 2019-nCoV_N2 Probe 5'-FAM-ACAATTTG CCCCCAGCGCTTCAG-BHQ1-3'). 18S rRNA (eukaryotic 18S rRNA endogenous control; Applied Biosystems) was used as an internal control to confirm appropriate specimen collection. The 18S rRNA probes were tagged with a 5'-VIC fluorophore and 3'-TAMRA quencher for multiplexing. Positive-sense vRNA for the standard curve was developed in-house by *in vitro* transcription, using the mMessage mMachine T7 kit (Ambion) and following the manufacturer's instructions. The limit of detection (LOD) for each one-step qPCR reaction was 23.2 genome copies. The final LOD based on 1 ml or 100 mg of sample was 1,856 genome copies/ml or 100 mg of tissue.

**Serology**. Serum neutralizing capacity was determined using an 80% plaque reduction neutralization test ($PRNT_{80}$) as described (*1*). Virus-specific total IgG and IgM were measured using ELISAs. ELISA plates (Maxisorp) were coated with 50 ng/well of SARS-CoV-2 RBD (kindly provided by Dr. Seema Lakdawala and prepared according to (*6*)) diluted in PBS overnight at 4 ºC. Plates were blocked in 5% (v/v) FBS, 5% (w/v) skim milk in PBS with 0.1% (v/v) Tween-20 for 1 hour at 37 ºC. Serial dilutions of plasma were made in block and incubated on blocked plates for 2 h at 37 ºC. Three washes with PBST were performed followed by incubation with goat-anti-monkey IgM(μ)-HRP (Seracare/KPL # 5220-0334) or goat-anti-rhesus IgG (H+L)-HRP (Southern Biotech # 6200-05), both used at a 1:5,000 dilution in blocking solution for 1 hour at 37 ºC. Three washes with PBST were performed prior to assay development by incubation with TMB (Seracare) for 7 min prior to the addition of TMB stop solution (Seracare). Absorbance values were determined at 450 nm.

**Pathology**. Tissues were immersion-fixed in 4% (w/v) paraformaldehyde, routinely processed, cut into 4 μm slides, and stained with hematoxylin and eosin (H&E). Stained slides were interpreted by a board-certified veterinary pathologist.

**Table S1. African green monkey cohort description**

| Animal ID | Sex | Age (years) | Weight (kg) | Infection Route | Exposure dose ($log_{10}$ pfu) | Euthanasia dpi | Outcome |
|---|---|---|---|---|---|---|---|
| A1 | M | 3.7 | 5.3 | Aero | 4.2 | 35 | mild disease |
| A2 | M | 3.6 | 6.3 | Aero | 4.0 | 35 | mild disease |
| A3 | M | 3.6 | 4.9 | Aero | 4.2 | 28 | mild disease; infection at telemetry implant site |
| A4 | M | 3.5 | 4.4 | Aero | 3.7 | 28 | mild disease |
| M1 | M | 3.3 | 4.2 | Multi-route mucosal | 6.4 | 28 | mild disease |
| M2 | M | 3.2 | 4.3 | Multi-route mucosal | 6.4 | 28 | mild disease |

**Table S2: Markers used for lymphoid and myeloid flow cytometry panels**

| Lymphoid | Myeloid |
|---|---|
| CD14, CD16 (dump) | CD3, CD20, NKG2A (dump) |
| Live/dead | Live/dead |
| CD3 | CD14 |
| CD4 | CD16 |
| CD8 | CD11c |
| HLA-DR | HLA-DR |
| NKG2A | CD123 |
| CD38 | CD38 |
| CD27 | Ki-67 |
| CD20 | |
| Ki-67 | |

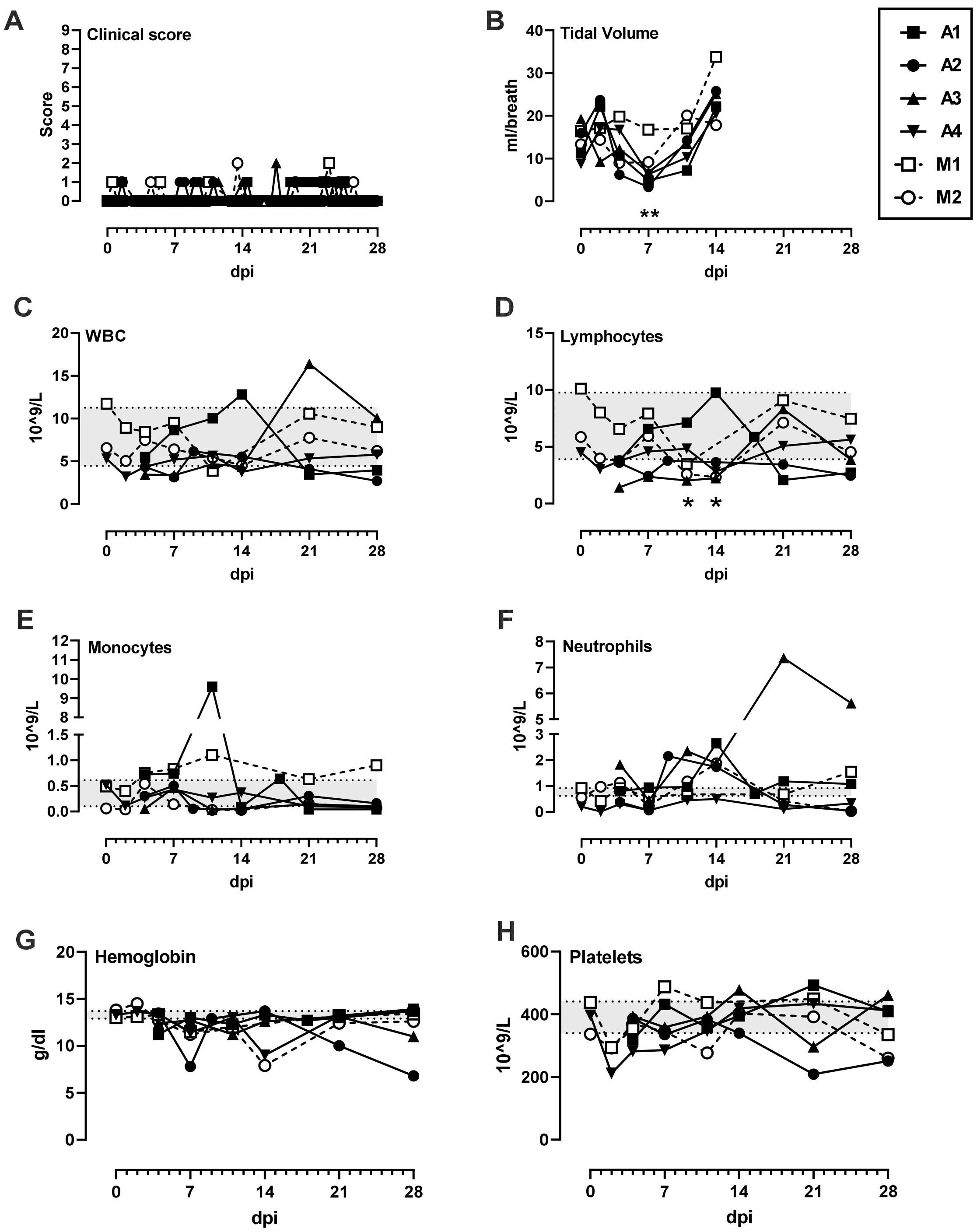


**Fig. S1. Clinical parameters in SARS-CoV-2 infected AGM**. (A) Total clinical score based on a summation of temperature, appearance/behavior, and respiratory signs (maximum score = 9). (B) Tidal volume measured by plethysmography. (C) whole blood cell count (WBC), (D) lymphocytes, (E) monocytes, (F) neutrophils, (G) hemoglobin, and (H) platelets. Statistical significance determined by ANOVA using GraphPad Prism and indicated by asterisks. Gray shaded area in C-F represent the mean pre-infection level +/- 1 standard deviation. AGM infected by aerosol (closed symbols/solid lines; n=4) or multi-route mucosal (open symbols/dashed lines; n=2).

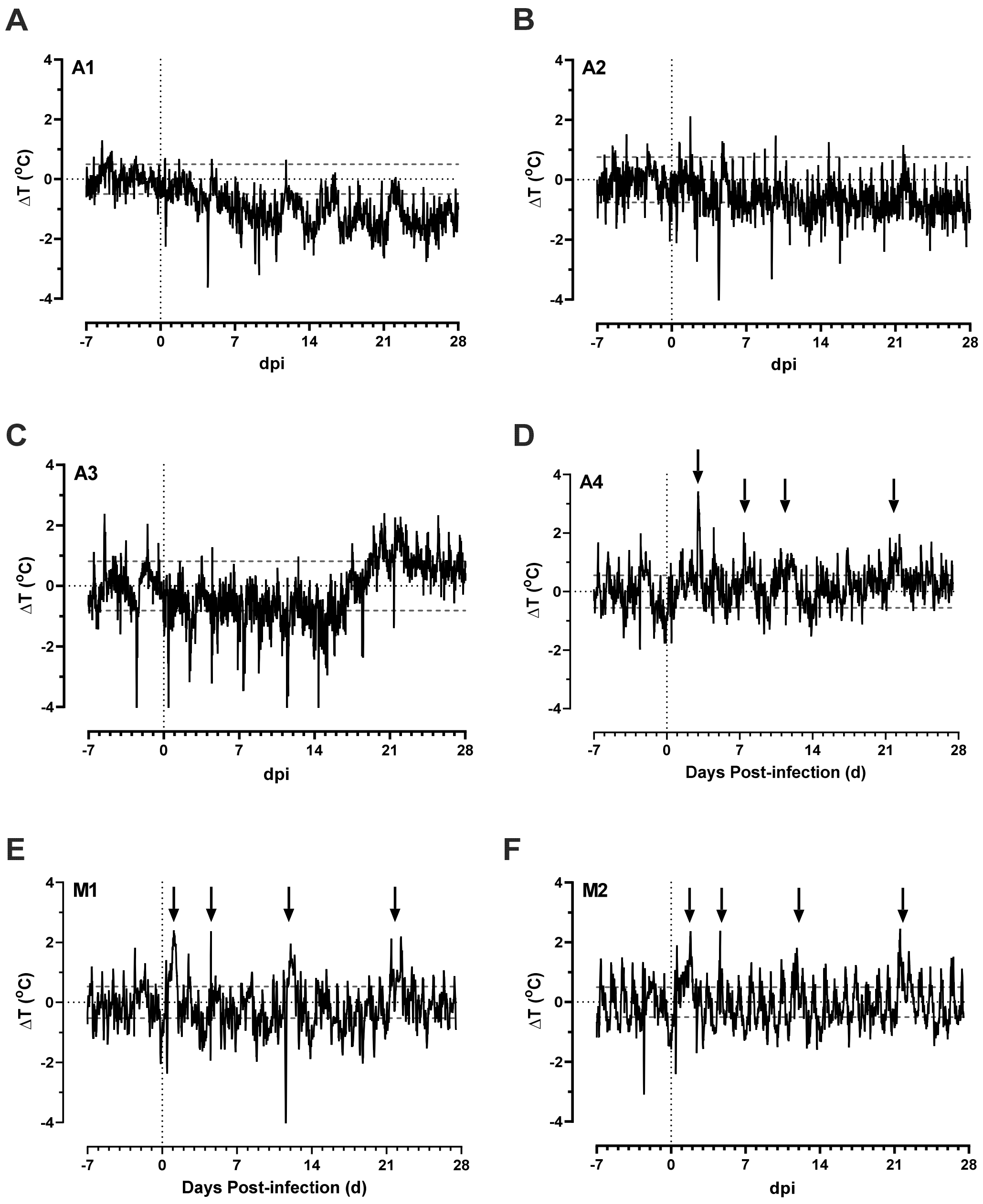


**Fig. S2. Body temperature changes in SARS-CoV-2-infected AGM.** Change in temperature over baseline for indicated animals. Horizontal dashed lines represent the upper and lower limit based on pre-infection data for each animal. Arrows indicate significant elevations.

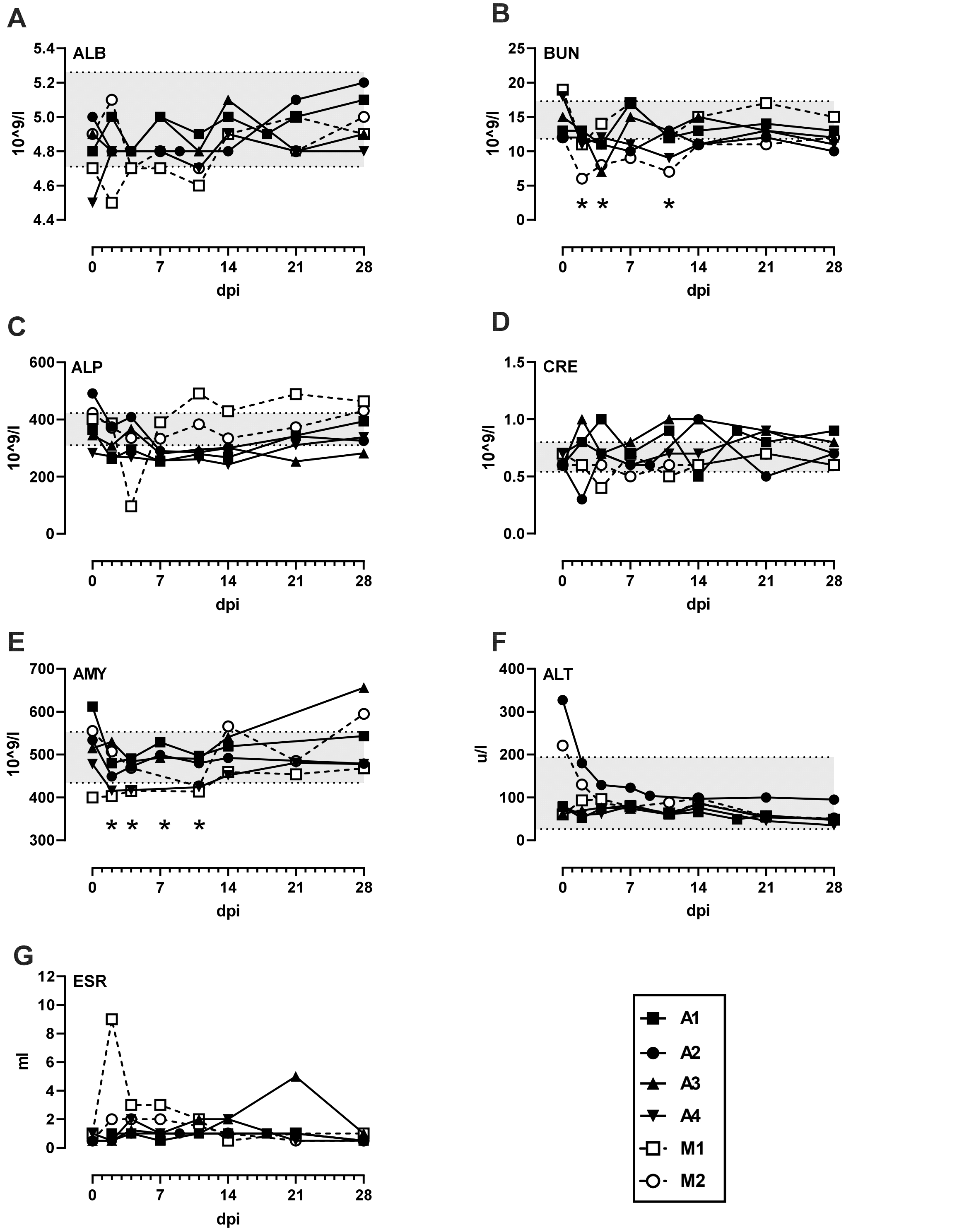


**Fig. S3. Blood chemistry parameters during SARS-CoV-2 infection**. (A) albumin, (B) blood urea nitrogen, (C) alkaline phosphatase, (D) creatinine, (E) amylase, (F) alanine transaminase, (G) erythrocyte sedimentation rate (ESR). Statistical significance was determined by 2-way ANOVA with multiple comparisons. Asterisks indicate significant changes compared to baseline (time 0) for each animal compared to its own baseline. Gray area in graphs A-F represent the mean of the pre-infection values +/- 1 standard deviation.

**(A) AGM A1**

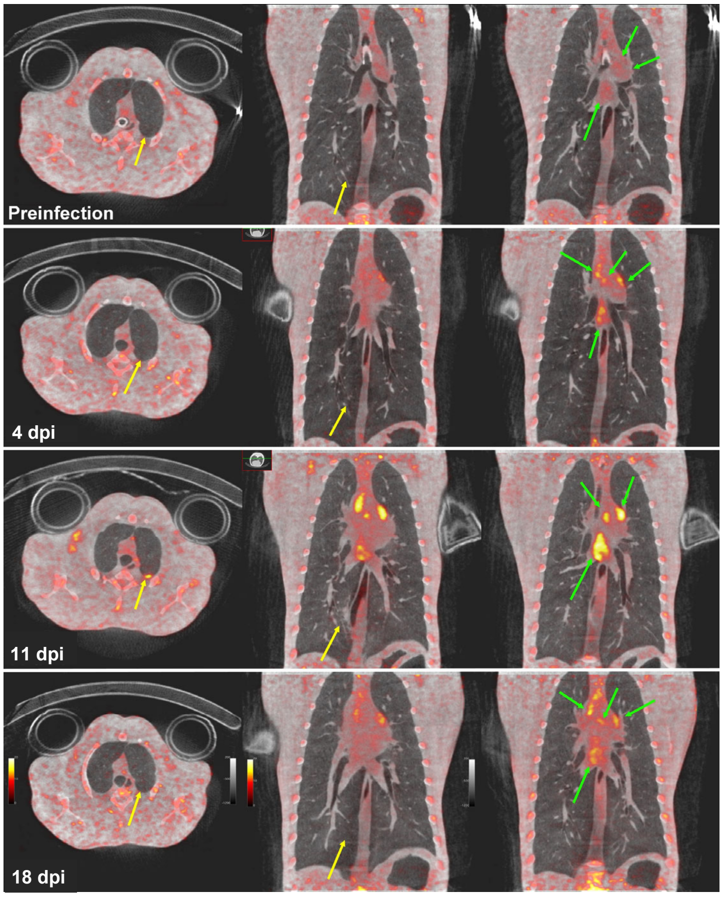


**(B) AGM A2**

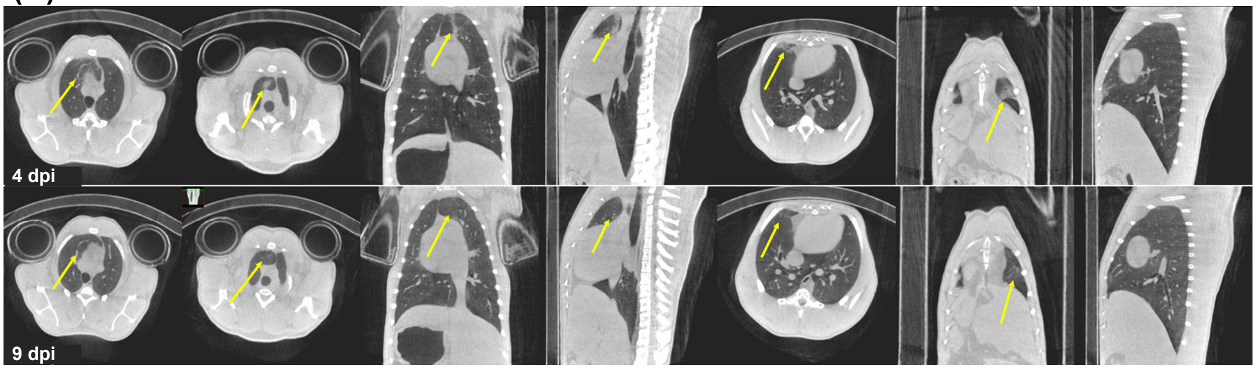


**Fig. S4. PET/CT images of AGM A1 and A2 infected with SARS-CoV-2**.

(A) AGM A1 (aerosol). PET/CT scans were obtained pre, 4, 11, and 18 dpi. No abnormal lung tissue at 4 dpi but the LNs were FDG avid. A focal pleural lesion was observed at 11 dpi on the posterior surface of the left upper lobe (FDG+) and an area of opacity in the medial portion of the right lower lobe (FDG-). By 18 dpi, lesions were resolving. Pulmonary infection (yellow arrows); thoracic lymph (green arrows). PET color scale is from 0 to 15 SUV.

(B) AGM A2 (aerosol). Only CT scans were obtained at 4 and 9 dpi. On day 4 dpi, A2 had ground glass opacity and thickened vessel structures in the anterior portions of the right upper and right middle lobes and a dense area of disease running vertically through the anterior portion of the right upper lobe.  On 9 dpi, some slight opacity was still present in the right middle lobe, but there was no abnormality in the right upper lobe.

**(A) AGM A3**

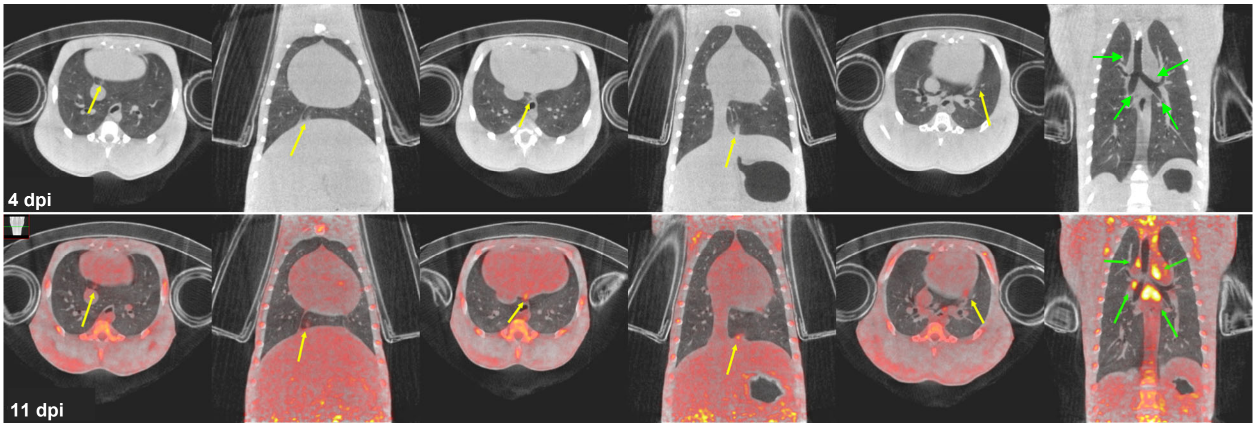


**(B) AGM M2**

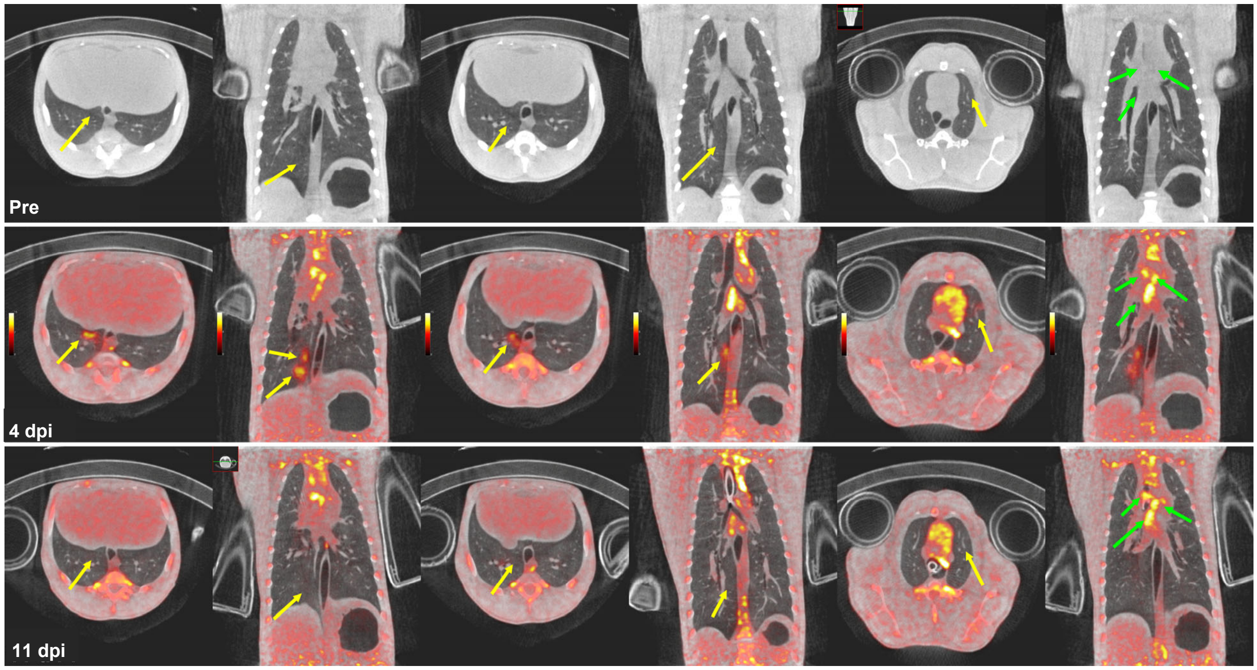


**Fig. S5. PET/CT images of AGM A3 and M2 infected with SARS-CoV-2**.

(A) AGM A3 (aerosol). A CT scan was obtained at 4 dpi and PET/CT at 11 dpi. At 4 dpi, two foci were visualized in the accessory lobe that were also present at 11 dpi (FDG+).

(B) AGM M2 (mucosal). An area of ground glass opacity, a distinct linear-shaped dense lesion in the right lower lobe, and an area of mid-parenchymal opacity in the left upper lobe. All were FDG+ and resolved by 11 dpi. Lymph node FDG uptake was consistent between 4 and 11 dpi. Pulmonary infection (yellow arrows); thoracic lymph nodes (green arrows). PET color scale is from 0 - 15 SUV.

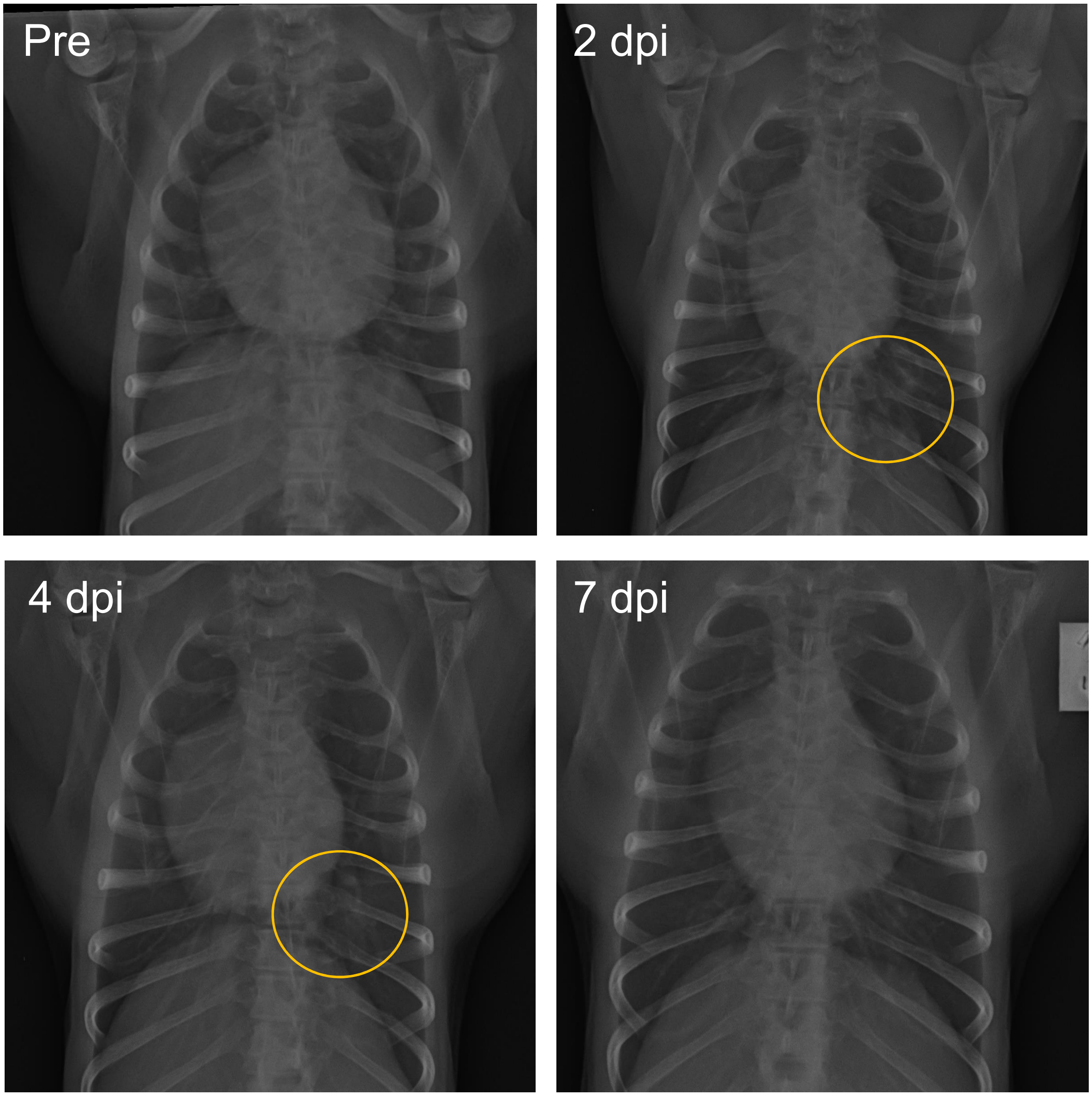


**Fig. S6. Mild lung infiltrates in SARS-CoV-2 infected AGM visualized by radiography**. Radiographs from AGM M1 (mucosal) at pre, 2, 4, and 7 dpi. Mild non-specific infiltrates were seen in the left lower lobe at 2 and 4 dpi that is resolving by 7 dpi. Radiographs were interpreted by a board-certified radiologist.

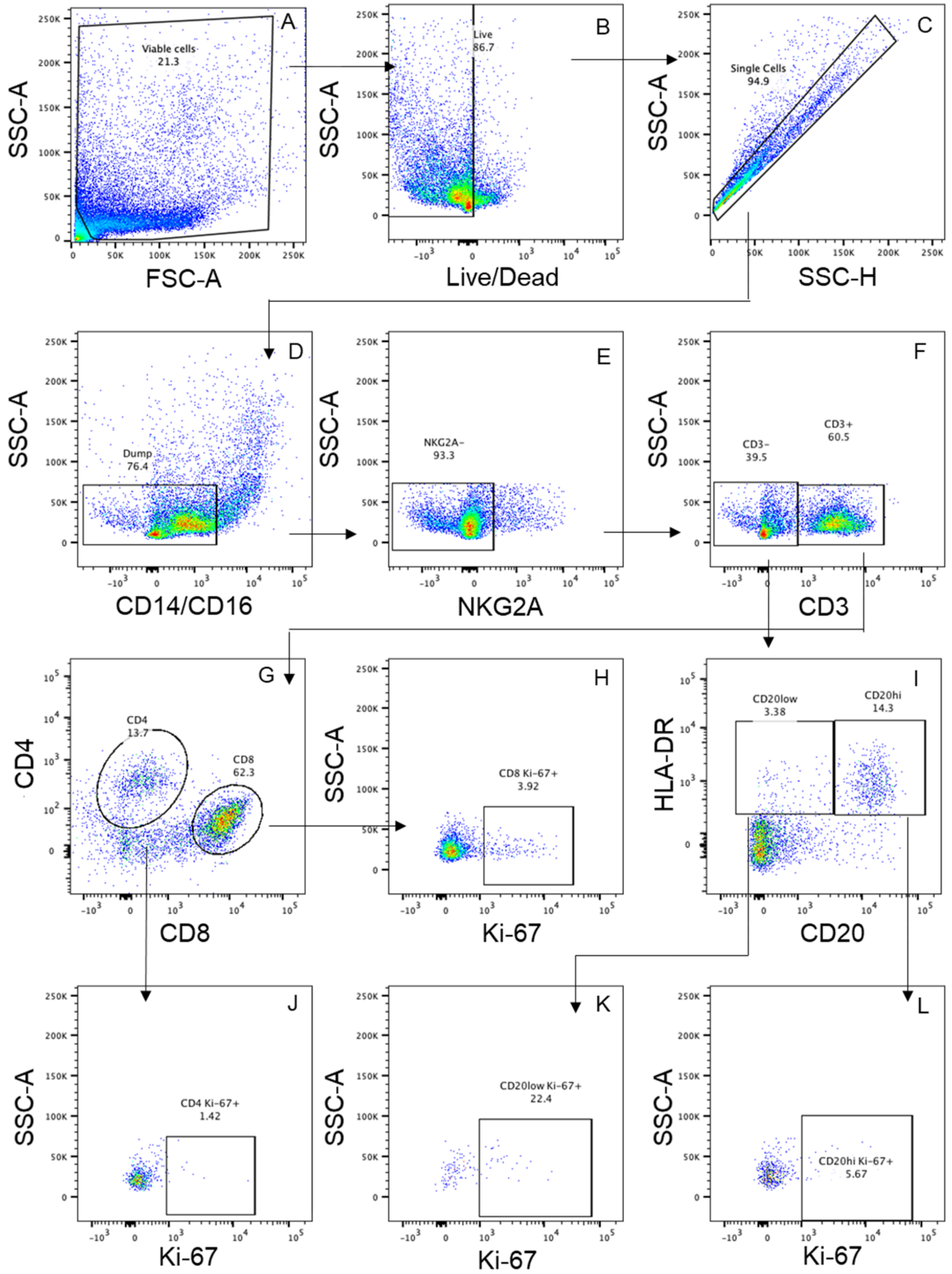


**Fig. S7: Lymphoid Panel Gating Strategy**. Representative PBMC sample from 0 dpi. Gating began with collection gate (A), live/dead (B), and singlet inclusion (C). CD14 and CD16 (D) and NKG2A (E) events were excluded from analysis and CD3 expression was evaluated (F). CD3+ events were further separated in CD4 or CD8 (G). CD4+ and CD8+ were then characterized by Ki-67+ (J and H, respectively). CD3- events were evaluated for CD20hi/low HLA-DR+ (I) CD20hiHLA-DR+ and CD20lowHLA-DR+ events were characterized for Ki-67+ (K and L).

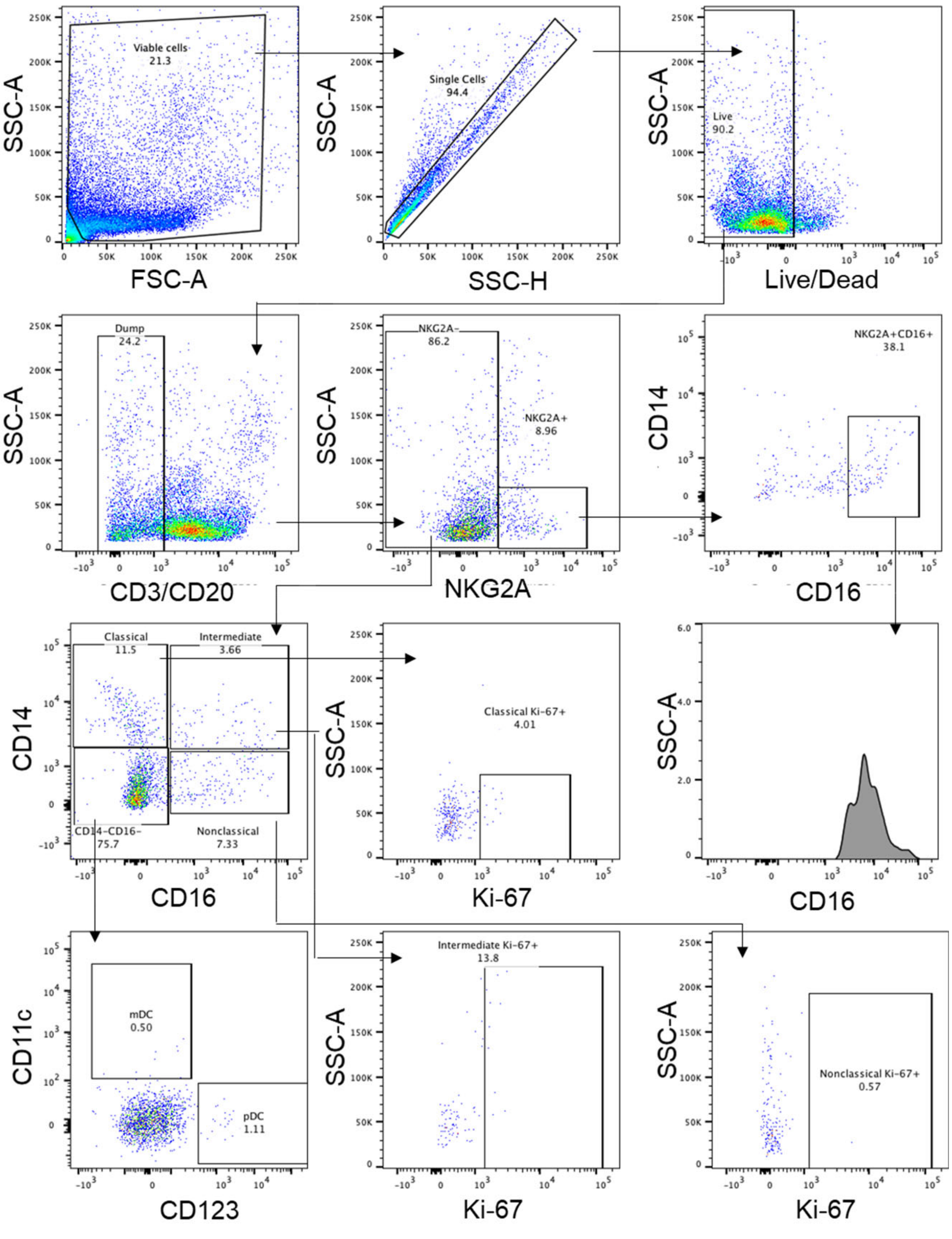


**Fig. S8: Meyloid Panel Gating Strategy**. Representative PBMC sample from 0 dpi. Gating began with viable cell gate (A), singlet inclusion (B), and Live/Dead (C) (same graphs as supplemental figure 5). CD3, CD20 (D) and NKG2A (E) were then excluded from analysis and evaluated by CD14 and CD16. CD14+CD16-, CD14+CD16+, and CD14-CD16+ were classified as classical, inflammatory, and nonclassical monocytes respectively (G). Classical (I and L), inflammatory (H and F), and nonclassical (K and M) were characterized for CD38+ and Ki-67+, respectively. CD14- events were evaluated for CD1c (mDC) or CD123 (pDC) (J).

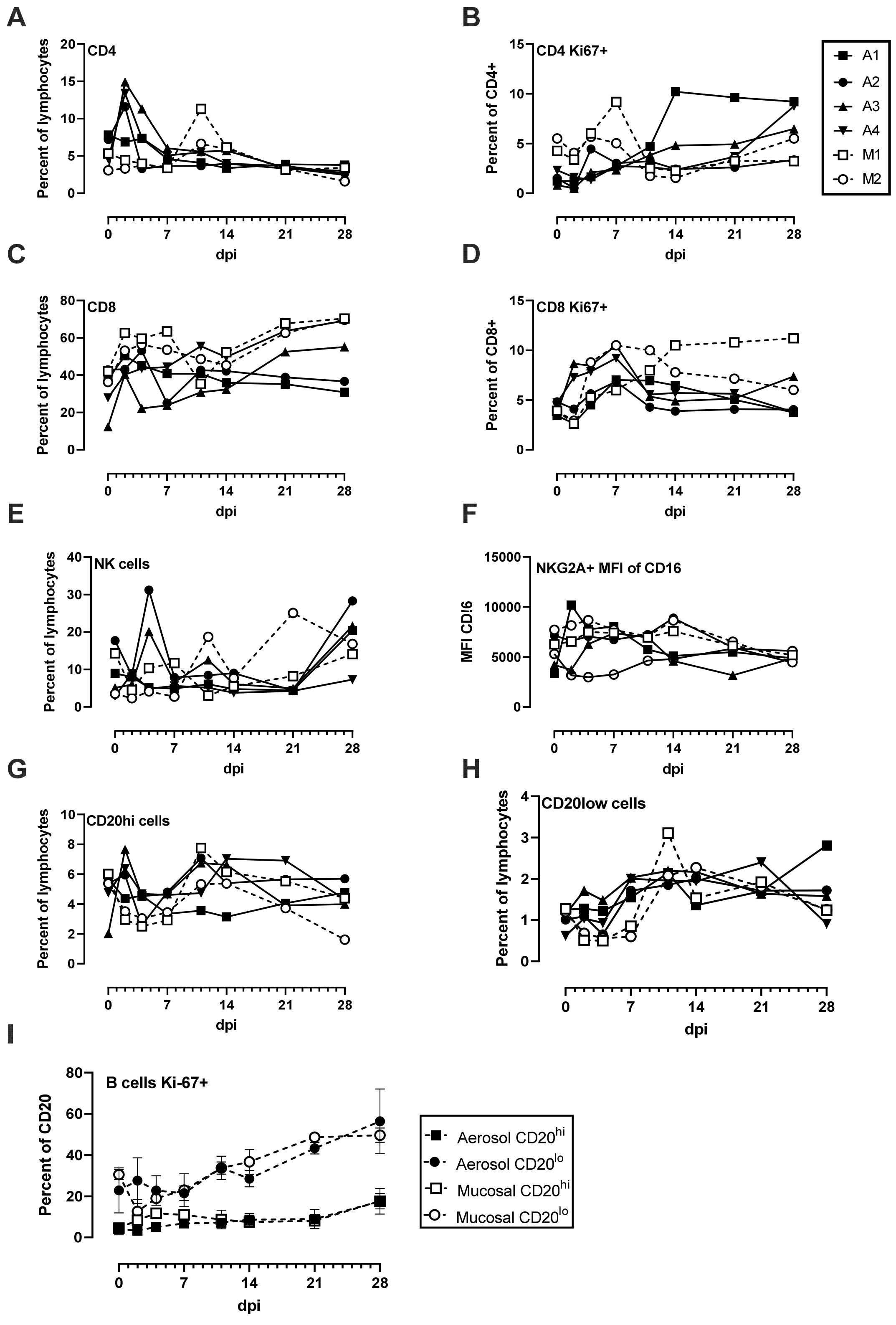


**Fig. S9: Lymphoid cell populations in SARS-CoV-2 infected AGMs**. (A) CD4+ T cells, (B) CD4+Ki-67+ cells, (C) CD8+ T cells, (D) CD8+Ki-67+ cells, (E) NK cells, (F) mean fluorescence intensity (MFI) of CD16 expression on NK cells, (G) B cells with high CD20 expression, (H) B cells with low CD20 expression (plasmablasts), (I) Grouped expression of Ki-67 on $CD20^{hi}$ and $CD20^{lo}$ (plasmablast) populations.

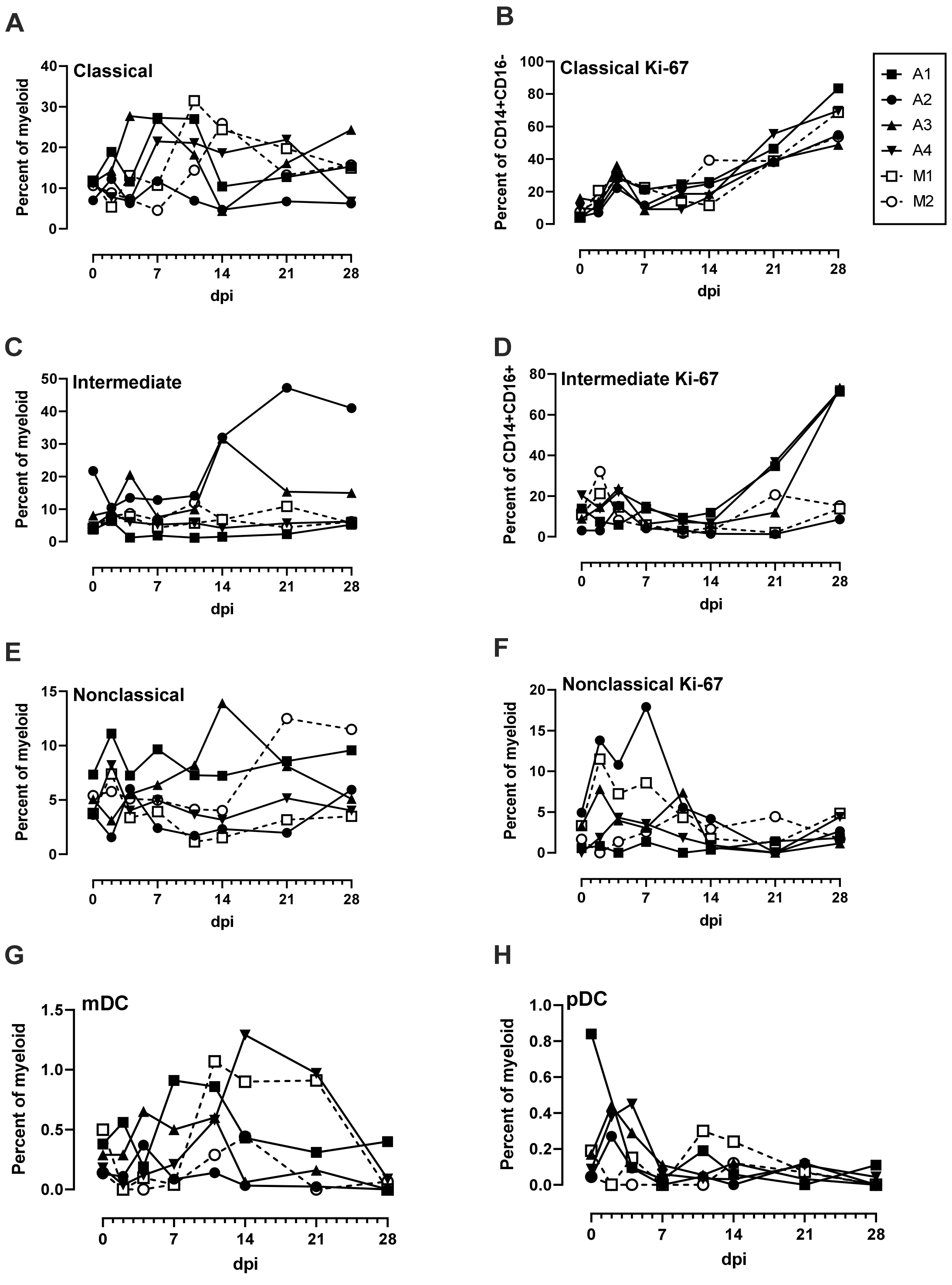


**Fig. S10: Monocyte populations in peripheral blood of SARS-CoV-2 infected AGMs**. (A) Classical monocytes (CD14+CD16-), (B) Ki-16+ Classical monocytes, (C) inflammatory monocytes (CD14+CD16+), (D) Ki-67+ inflammatory monocytes, (E) nonclassical monocytes (CD14-CD16+), (F) Ki-67+ nonclassical monocytes, (G) myeloid dendritic cells (mDCs), and (H) plasmacytoid dendritic cells (pDCs).

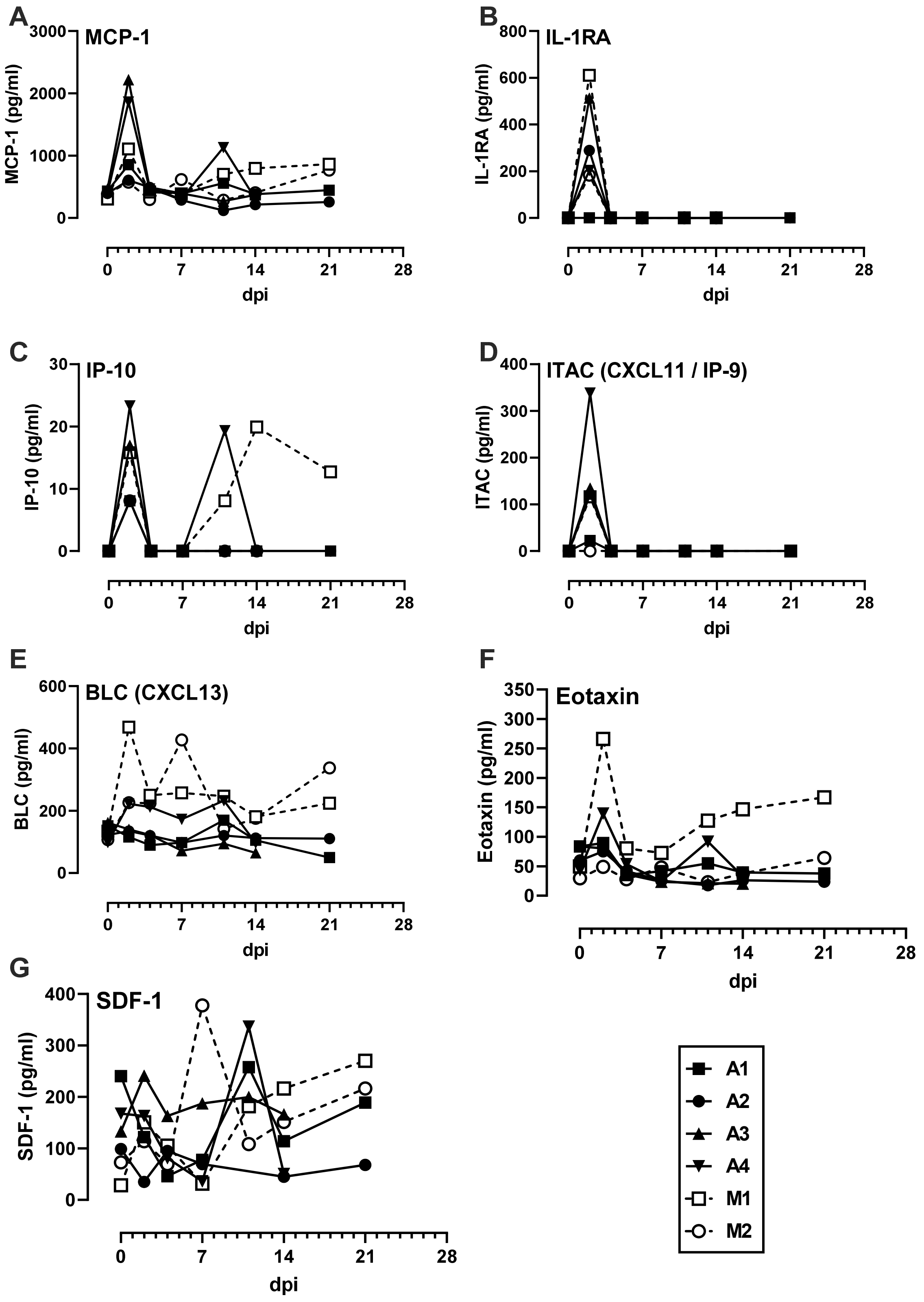


**Fig. S11. Cytokinemia during SARS-CoV-2 infection**. The indicated parameters were measured in longitudinal plasma samples using the Cytokine & Chemokine 30-Plex NHP ProcartaPlex Panel from Invitrogen. Parameters not shown were below the limit of detection across all animals and time points. Aerosol (closed symbols/solid lines; n=4); multi-route mucosal (open symbols/dashed lines; n=2).
